# S-Nitrosylated COX-2 is a TME-regulated breast cancer biomarker of mesenchymal phenotypes

**DOI:** 10.1101/2025.07.15.664474

**Authors:** Reuben J Hoffmann, AeSoon Bensen, Mark Dane, Jane Arterberry, Rebecca Smith, James Korkola, Pepper Schedin

## Abstract

COX-2 is an inducible enzyme key to the production of inflammatory prostaglandins. COX-2 also has tumor intrinsic oncogenic activity in mouse models of breast cancer. Previously, we reported increased expression of Cys-526-nitrosylated COX-2 (SNO-COX-2), but not non-nitrosylated COX-2, with progression of early-stage human breast cancer to invasive ductal carcinoma. Here, we used a 3D culture model of early-stage human breast cancer (MCF10DCIS cells) to investigate the relationship between SNO-COX-2 expression and mesenchymal/invasive tumor cell morphology. We find that SNO-COX-2, but not non-nitrosylated COX-2, closely associated with mesenchymal phenotypes induced by fibrillar type I collagen. Interestingly, invasive phenotypes did not associate with induction of the classic epithelial-to-mesenchymal transition (EMT) markers *SNAIL*, *CDH2* (N-cadherin), and *VIM* (vimentin). By contrast TGFβ-1 strongly induced EMT-related transcripts, but not SNO-COX-2 protein expression or mesenchymal phenotypes. These observations suggest that in MCF10DCIS cells, SNO-COX-2 associates with mesenchymal phenotypes more strongly than non-nitrosylated COX-2 protein, or expression of classic EMT transcripts. In a mouse model with breast tumor heterogeneity, mesenchymal tumor regions also have increased SNO-COX-2 expression. Testing 300 distinct tumor microenvironment conditions, we find SNO-COX-2 protein expression is driven by inflammation, wound resolution, and cancer-associated factors, especially TNC, SPP1, decorin, fibrillar type I and III collagens, INF-γ, and IL-4/13, with evidence for specific extracellular matrix-ligand interactions driving both high and low SNO-COX-2 expression. In sum, in MCF10DCIS cells, expression of SNO-COX-2 is highly microenvironment-dependent and strongly associated with invasive/mesenchymal growth, indicating potential for SNO-COX-2 as a biomarker to assess risk of early-stage breast cancer progression.

## Introduction

Inflammation is a hallmark of cancer during both early- and late-stage disease progression [1, 2]. The enzyme cyclooxygenase-2 (COX-2, PTGS2) produces the major precursor of the prostanoids required for production of inflammatory prostaglandins. *COX-2* gene expression is induced by various inflammatory conditions, including cancer [3]. High levels of COX-2 and its downstream products are linked to increased cancer incidence and worse prognosis in many solid tumors including breast [4], prostate [5], and colon [6]. COX-2 has also been shown to act as an oncogene in a rodent model of breast cancer [7]. Inhibition of COX-2 by non-selective and selective non-steroidal anti-inflammatory drugs (NSAIDs), including aspirin, ibuprofen, and celecoxib, is sufficient to reduce inflammation [8], and NSAID treatment has preventative and therapeutic activity in various rodent cancer models including breast [9, 10]. Further, a recent breast cancer study found that regular post-diagnostic aspirin use associated with a 38% and 28% reduced risk of breast cancer deaths [11]. As such, COX-2 signaling remains an important pro-tumor pathway with continued promise for drug development. However, barriers remain with respect to the effective targeting of COX-2 for the prevention or treatment of breast cancer, including the pleiotropic and opposing nature of COX-2 downstream products depending on cell type and localization [3] and evidence for post-translational COX-2 protein modifications with implications for protein function [12].

COX-2 has been shown to be post-translationally modified on Cystine-526 by S-nitrosylation, a common yet understudied protein modification in which a nitro group (-NO) is added to the thiol group of a cystine (-SH), to form a nitrosothiol group (-SNO). S-nitrosylation of COX-2 has been demonstrated to increase COX-2 enzymatic activity *in vivo*, in cells, and in lysate [13]. One source of cellular nitric oxide (NO) is inducible nitric oxide synthase (iNOS; NOS2). iNOS and COX-2 protein levels are frequently correlated, implicating similar inflammatory control. Like COX-2, iNOS is involved in downstream inflammatory signaling via its product, NO [13]. High iNOS expression is associated with worse prognosis in breast cancer and shows promise as a therapeutic target [14, 15], yet specific nitrosylated protein targets are largely understudied, including COX-2.

The Schedin lab has identified and validated commercially available antibodies that delineate between the cystine-526-S-nitrosylated and cystine-526-non-nitrosylated forms of human COX-2 [16], hereafter **SNO-COX-2** and **n-COX-2**, respectively. In prior work, utilizing these antibodies to stain normal human breast tissue, it was found that the nitrosylated and non-nitrosylated forms of COX-2 have distinct regulation and intracellular localizations, implicating distinct functions [16]. Further, in a small cohort of breast cancer cases that contained far normal, adjacent normal, ductal carcinoma *in situ* (DCIS), and invasive breast cancer on the same tissue section, increased SNO-COX-2, but not n-COX-2, associated with tumor progression [16]. As additional support for SNO-COX-2 being a potentially improved biomarker of COX-2 oncogenic activity, SNO-COX-2 expression is prevalent in poor prognostic young women’s breast cancer compared to breast cancers diagnosed in postmenopausal women [16] and associates with normal weaning-induced breast involution, a reproductive window of vulnerability for young onset breast cancer [17, 18]. Combined, these observations indicate previously unrecognized functional differences for SNO-COX-2 and n-COX-2 in normal mammary epithelial cells and implicate SNO-COX-2 as particularly relevant to breast cancer progression.

Here, using a previously described 3D cell culture model of early-stage breast cancer, we investigated associations between S-nitrosylation of COX-2 and tumor cell invasive attributes. The human breast cancer cell line MCF10DCIS.com (hereafter MCF10DCIS) forms non-invasive DCIS-like lesions *in vivo* that progress to invasive disease with time, making this cell line a useful model of early-stage disease [19–21]. In 3D cell culture, MCF10DCIS cells form non-invasive DCIS-like spheroids when grown in a laminin 1 enriched microenvironment (i.e., Matrigel) but can be induced into invasive growth by the addition of fibrillar type I collagen [22, 23]. *In vivo* and in cell culture models, tumor cell invasion is frequently driven by epithelial-to-mesenchymal transition (EMT) programs [24]. EMT is a dynamic morphological transition that leads to tumor cells with mesenchymal morphology and increased motility, invasiveness, and metastatic potential [24]. In many cancer cells, EMT is induced by COX-2, type I collagen and TGFβ-1 [20, 25]. Here, we investigated associations between SNO-COX-2 protein levels, tumor cell mesenchymal morphology, and EMT biomarkers in 3D culture. We also interrogated over 300 distinct tumor microenvironment (TME) conditions in 2D culture. Combined our studies more fully describe extrinsic mediators of SNO-COX-2 expression and associated mesenchymal phenotypes in MCF10DCIS cells.

## Results

### SNO-COX-2 is associated with EMT-like morphology

Our model consists of MCF10DCIS, a c-Ha-ras oncogene-transformed derivative of the immortalized human non-cancer MCF10A breast epithelial line [19, 20], grown under distinct TME conditions. A DCIS-like breast cancer model is ideal for exploring potential roles for SNO-COX-2 in disease progression, as the DCIS to IDC transition is thought to be largely TME-mediated [20, 23]. MCF10DCIS cells were plated into 3D culture pads consisting of Matrigel with or without the addition of fibrillar type I collagen (hereafter +Col1 or +C), as an established pro-tumor, pro-EMT, extracellular matrix (ECM) protein [22, 23, 25]. After 24 hours, a timeframe that supported expansion of single cells into small (2-3 cell), compact tumor spheroids (**Sup. Fig. 1A**), the cultures were treated with control diluent or EMT-inducer TGFβ-1 [25, 26] (hereafter +TGFB or +T). Bright field imaging confirmed equal cell density at time of plating across conditions and captured spheroid growth and morphologic changes over time (**Fig 1A, Sup. Fig. 1A**). At 5- or 7-day study endpoints, Matrigel pads were processed for RNA isolation or fixed for subsequent immunohistochemical analyses. Because MCF10DCIS is known to be bipotent and capable of forming DCIS-like lesions surrounded by myoepithelial-like cells, we stained our 3D Matrigel cultures for the myoepithelial cell marker p63. We found p63 expression in spheroid basal cells, as defined by cells in direct contact with Matrigel (**Sup. Fig. 1B**), consistent with acinar formation, the dual epithelial/myoepithelial character of MCF10DCIS cells, and utility of this cell line for the study of early-stage breast cancer[19, 20].

**Figure 1:**
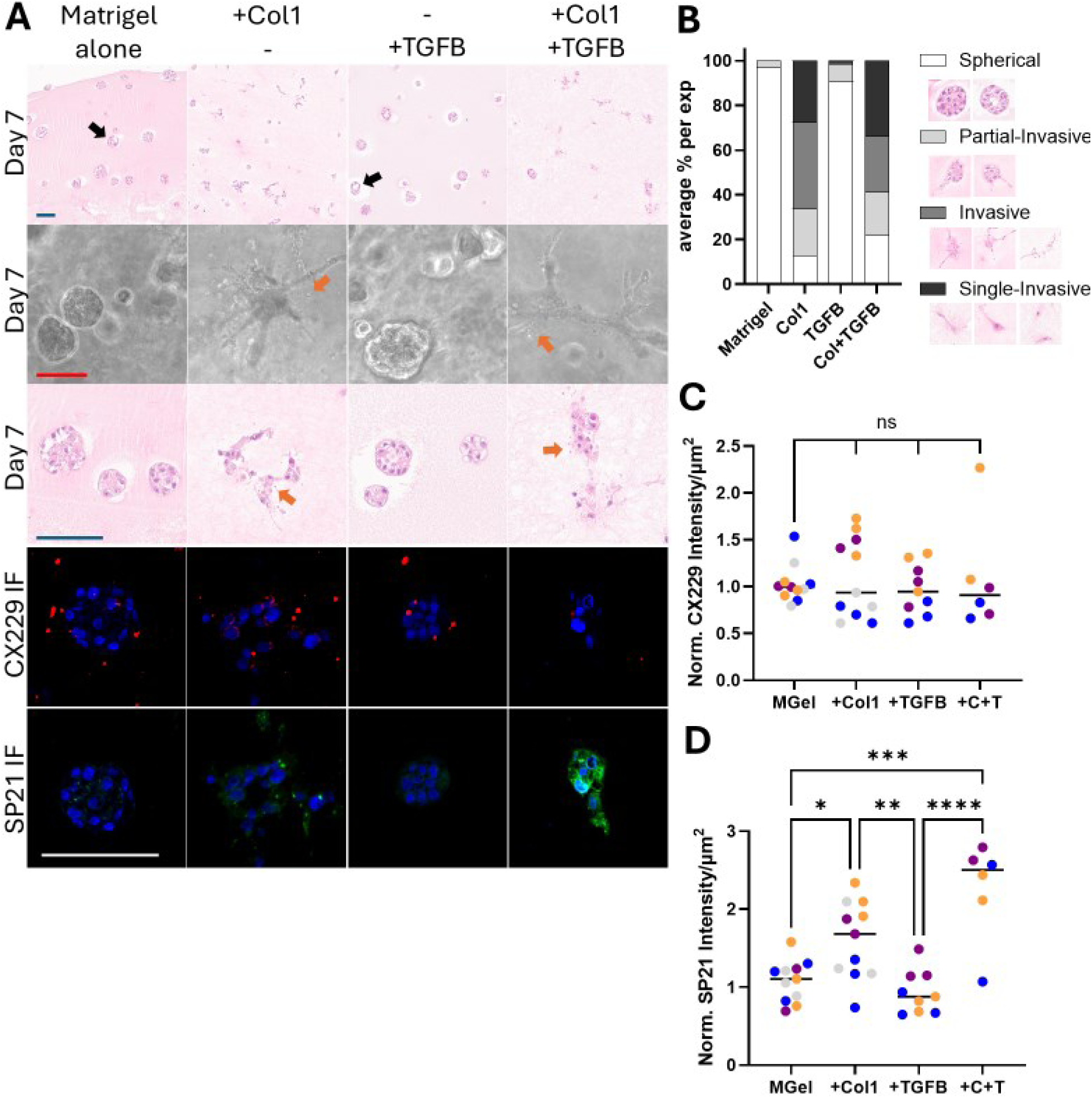
Effects of fibrillar type I collagen and TGFβ-1 on mesenchymal phenotypes and SNO-COX-2 expression. **A)** MCF10DCIS.com cells grown in 3D Matrigel culture with and without fibrillar type I collagen and TGFβ-1. All images taken from day 7 cultures. Upper panel, low magnification images of sectioned FFPE spheroids H&E stained, black arrows identify tumor spheroids with acinar-like structures. Second panel shows high magnification bright field images, and third panel shows high magnification sectioned FFPE spheroids H&E stained. Orange arrows depict mesenchymal phenotypes. Lower two panels show representative IF staining for n-COX-2 (red) and SNO-COX-2 (green), with DAPI stained nuclei (blue). All scale bars = 100 µm. **B)** Quantitation of spheroid morphology by category: spherical, partially invasive, multicellular invasive, and single-cell invasive, with representative images for each phenotype shown in side panel. Events were counted in 3-4 images from each condition in triplicate across four experimental replicates. The average percent of each phenotype in each ECM condition is shown as a stacked bar graph. +C and +C+T conditions were significantly different from control in all phenotypes, details in Sup. Fig. 1D. **C)** Quantitation of n-COX-2 (CX229) IF staining across culture conditions. Colors indicate distinct experiments. Individual points represent the mean of each technical triplicate. Black bars indicate mean-of-means from all replicates. **D)** As C, for SNO-COX-2 (SP21) IF staining. Significance is shown as *, **, *** and **** for p values < 0.05, 0.01, 0.001, and 0.0001, respectively. “ns” or unlisted relationships were not significant (α = 0.05)

To quantify morphological changes between the distinct TME conditions – Matrigel alone, +Col1, +TGFB, and +Col1+TGFB – we binned spheroid and cell morphologies into four morphological categories: 1) round, smooth edged spheroids 2) partially invasive spheroids (overall round with few cellular extensions/protrusions) 3) invasive (multicellular with frequent cellular extensions/protrusions) 4) and single invasive cells (elongated single cells consistent with full morphologic EMT). Percentage spheroid phenotypes per condition and examples of each phenotype are depicted in **Fig. 1B**. As expected, tumor spheroids in Matrigel were mostly round and smooth-edged (**Fig. 1A & B**). Somewhat surprisingly, given the established role of TGFβ-1 in inducing EMT in other breast cancer cell lines [25, 26], cells grown in Matrigel +TGFB formed round and smooth-edged spheroids (**Fig. 1A & B, Sup. Fig. 1A**). The Matrigel +TGFB condition spheroids were also slightly smaller in size than the Matrigel only condition (**Sup. Fig 1C**). Smaller size is consistent with the known antiproliferative effects of TGFβ-1 on normal epithelium and early stage disease [27], providing additional support for MCF10DCIS cells representing early-stage human breast cancer. Of note, no statistically significant difference between the percentage of any morphologic phenotype was observed between +TGFB and matrigel alone conditions (**Fig. 1A & B, Sup. Fig. 1D**). By contrast, MCF10DCIS cells cultured with fibrillar type I collagen, either alone or treated with TGFβ-1, led to a statistically significant gain of all three of the invasive phenotypes (∼20-30% gain each) and loss of rounded spheroids (∼75-85% loss) compared to the non-collagen conditions (**Fig. 1A & B, Sup. Fig. 1D**; all p <0.001 or <0.0001). Comparing the +Col1+TGFB condition to +Col1 alone, TGFβ-1 was associated with a small but significant increase in spherical events (**Sup. Fig. 1D**, ∼9% gain, p = 0.046) and a loss of mesenchymal events (**Sup. Fig. 1D**, ∼14% loss, p = 0.014), suggesting that TGFβ-1 is more inhibitory than EMT-inducing in MCF10DCIS cells grown in 3D culture.

We next addressed whether the invasive morphologic phenotypes observed in the type I collagen culture conditions associated with increased SNO-COX-2. To this end, we performed immunofluorescent (IF) staining for SNO-COX-2 and n-COX-2 using the previously validated n-COX-2-specific antibody clone CX229 (hereafter CX229) and the SNO-COX-2-specific antibody clone SP21 (hereafter SP21) [16]. We did not see changes in protein expression of n-COX-2 (CX229 staining) across the four different TME conditions tested, with equivalent, relatively low-level detection in non-invasive and invasive spheroids (**Fig. 1A, C**). However, we found SNO-COX-2 expression (SP21 staining) did correlate with invasive morphology, with low expression in both Matrigel-alone and Matrigel +TGFB conditions and significantly higher expression in the +Col1 and +Col1+TGFB conditions (**Fig. 1A, D**). While not reaching significance, the combination of fibrillar type I collagen and TGFβ-1 together trended towards higher SNO-COX-2 staining than the +Col1 condition alone (**Fig. 1A, D**). Ultimately, in this model system, SNO-COX-2 closely associated with invasive morphology induced by fibrillar type I collagen whereas n-COX-2 did not.

We observed that the invasive cellular phenotypes in the Col1+ conditions were accompanied by increased cellular debris, vesicles, and blebs, which were evident in both live (brightfield) and fixed cultures (**Fig. 1A & Sup. Fig. 1A; orange arrows**). To test whether these small cellular particles associated with a cell death phenotype, IHC staining for cleaved caspase 3 (CC3) was performed. In all conditions, CC3+ cells were rare, and were primarily found in spheroids that were hollowing (**Sup. Fig. 1E**), consistent with previously published studies demonstrating that acinar formation *in vivo* and *in vitro* can occur through a programmed cell death mechanism [28]. Quantification of CC3+ confirmed that the number of spheroids containing CC3+ cells was low and did not significantly differ between conditions (**Sup. Fig. 1F**). In sum, these data suggest that the increased SNO-COX-2 staining observed in the +Col1 conditions is not primarily driven by cell death but rather associates with the EMT-like phenotype in viable cells. Because COX-2 has been shown to be associated with EMT in other models and the invasive phenotype we observe resembles EMT, we next examine the expression of common biomarkers of EMT in our 3D model.

### Transcriptional expression of EMT-related genes does not associate with EMT morphology nor SNO-COX-2 expression

To determine if molecular biomarkers of EMT track with observed phenotypic EMT-like morphology and its associated SNO-COX-2 expression, RNA was isolated from MCF10DCIS cells after five days of 3D culture. The expression of three EMT-related genes, *SNAIL*, *VIM* (vimentin), and *CDH2* (*NCAD*; N-cadherin) were assessed by RT-qPCR. (**Fig. 2A-C**). SNAIL (SNAI1) is a key transcription factor that drives EMT [24]. Intriguingly, *SNAIL* expression was not upregulated by the +Col1 alone condition (**Fig 2A**). Similarly, vimentin and N-cadherin, downstream genes involved in intermediate filament reorganization associated with directional polarization (vimentin) and mesenchymal cell-cell adhesion (N-cadherin), were either not upregulated (vimentin) or only slightly upregulated (N-cadherin; 2-fold mean difference, p < 0.0001) in the +Col1 alone conditions (**Fig. 2B, C**). Conversely, *SNAIL*, vimentin and N-cadherin genes were strongly induced by +TGFB alone, from 2-to 44-fold mean difference (**Fig 2A-C**, p < 0.0001), yet TGFβ-1 did not induce morphological EMT (**Fig 1A-B**). In the combined +Col1+TGFB condition, *SNAIL* and N-cadherin expression was further increased about two-fold compared to +Col1 (**Fig 2A**, p < 0.0001), implicating a TGFβ-1 effect. It is notable that the observed increase in vimentin expression induced by TGFβ-1 was low when compared to SNAIL and N-cadherin, only about 2-fold (**Fig 2B**) compared to almost 10-fold and 44-fold increases for SNAIL or N-cadherin, respectively (**Fig 2A, C**). One explanation is that baseline expression of vimentin is high in MCF10DCIS cells, as demonstrated by low Cq values in the RT-qPCR data (**Sup. Table 1**) and high vimentin IHC staining in our 3D cultures (**Sup. Fig. 2A**).

**Figure 2:**
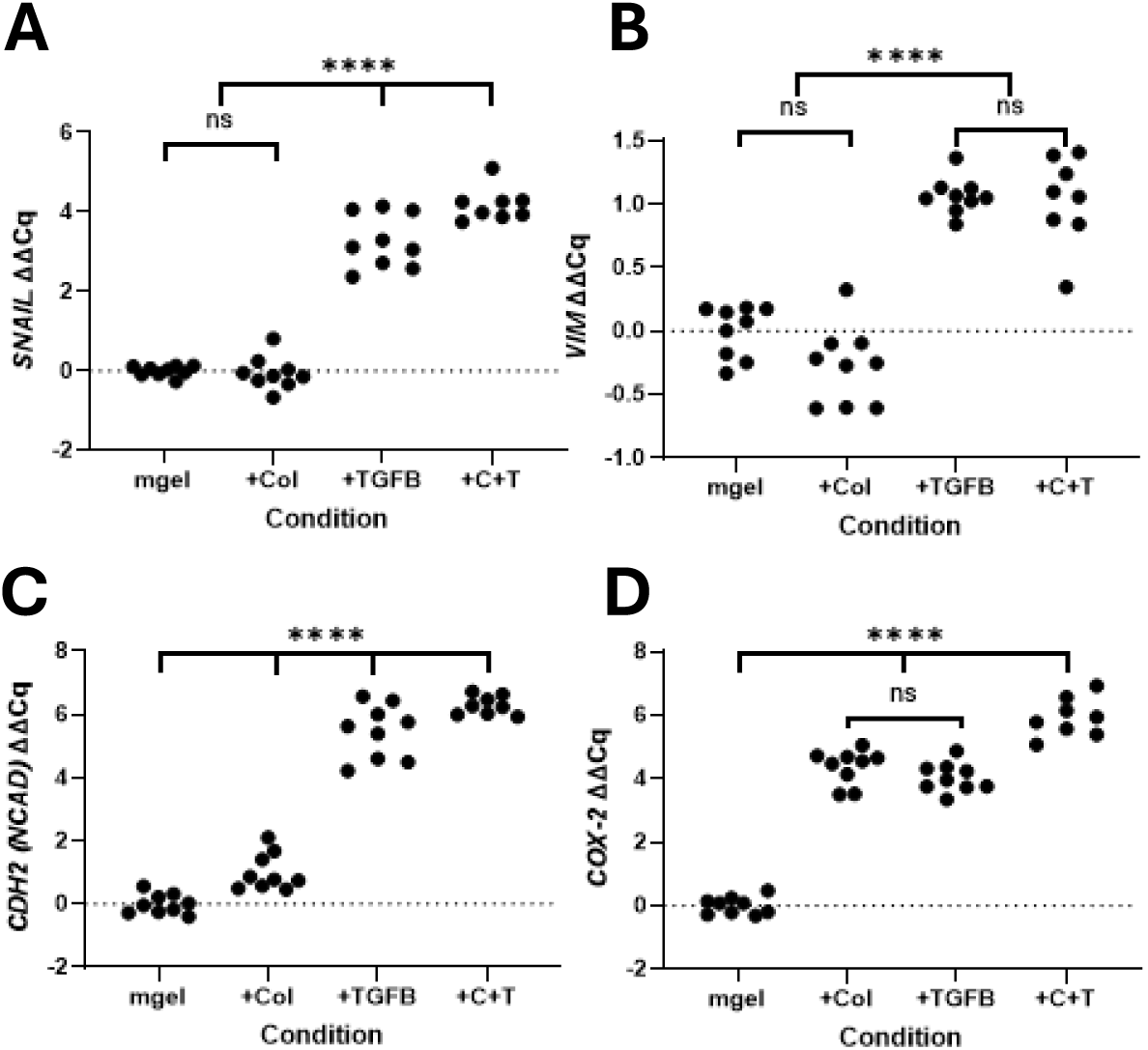
Gene expression of EMT-related biomarkers and COX-2. RT-qPCR was performed on RNA isolated from 3D cultures at Day-5. Results are shown for transcripts of **A)** *SNAIL*, **B)** Vimentin, **C)** N-cadherin, and **D)** *COX-2*, normalized against a GAPDH control in three experimental replicates performed in technical triplicate. The data are presented as ΔΔCq (equivalent to log_2_ of the fold change) calculated using the averages of the qPCR triplicates, with data normalized to matrigel control values. A lack of significant differences between two conditions is indicated by an “ns” bracket. All other condition-vs-condition differences are significant (p <0.001), indicated by an overarching **** bracket.

We next evaluated whether increased SNO-COX-2 protein correlated with increased *COX-2* gene expression. Both +Col1 and +TGFB induced *COX-2* gene expression to similar levels, 16-to 20-fold greater on average, while the combination of the two factors increased levels almost 60-fold over matrigel alone (**Fig 2D**, p < 0.0001). However, increased *COX-2* gene expression did not track with EMT/mesenchymal morphology; *COX-2* was 16-fold greater in the +TGFB condition compared to Matrigel despite the absence of mesenchymal morphology. These data demonstrate a disjunct between *COX-2* mRNA levels and COX-2 protein levels in MCF10DCIS cells, which is in line with previous studies reporting no correlation between *COX-2* RNA and protein levels by staining [29]. Because SNO-COX-2 has been previously reported to be influenced by iNOS levels [13], we also assayed *iNOS* (*NOS2*) transcription, as well as the *NOS* homologues *nNOS* (*NOS1*) and *eNOS* (*NOS3*). Curiously, we were unable to detect any nitric oxide synthase mRNA in our samples using five different sets of qPCR primers, all of which were functional in positive control RNA extracted from the acute myeloid leukemia line AML2 (**Sup. Fig. 2B**). Further, the addition of iNOS inhibitors L-NMMA and 1400W did not decrease SNO-COX-2 protein levels in our 3D model (data not shown). These data imply that the increase in SNO-COX-2 in our model system is independent of NOS activity.

In sum (**Table 1**), we had expected SNO-COX-2 expression and the EMT-like invasive phenotype induced by the +Col1 conditions to be correlated with the expression of classical EMT markers. Instead, we observed that increased transcription of the genes *SNAIL*, vimentin, and N-cadherin, as well as *COX-2* transcription itself was not sufficient to induce EMT-like invasive phenotypes (**Fig. 1 vs Fig. 2**, +TGFB condition). Further, we found that increased expression of *SNAIL* and vimentin were not required (necessary) for the EMT-like phenotype induced by +Col1 alone (**Fig 1 vs. Fig. 2A-B**, +Col1 condition). Of note, N-cadherin was marginally increased by +Col1 alone compared to +TGFB (**Fig. 2C**), thus we cannot rule-out the possibility that modest increases in N-cadherin expression are necessary for the EMT-like phenotypes observed. Overall, SNO-COX-2 protein level was the only tested factor to closely correlate with the observed EMT morphology (**Table1**). Because it is possible that SNO-COX-2’s association with EMT-like growth is dependent on a component unique to our model (cell line, matrigel culture, etc.), we next examined murine mammary tumors for an association between SNO-COX-2 and mesenchymal character.

**Table 1:**
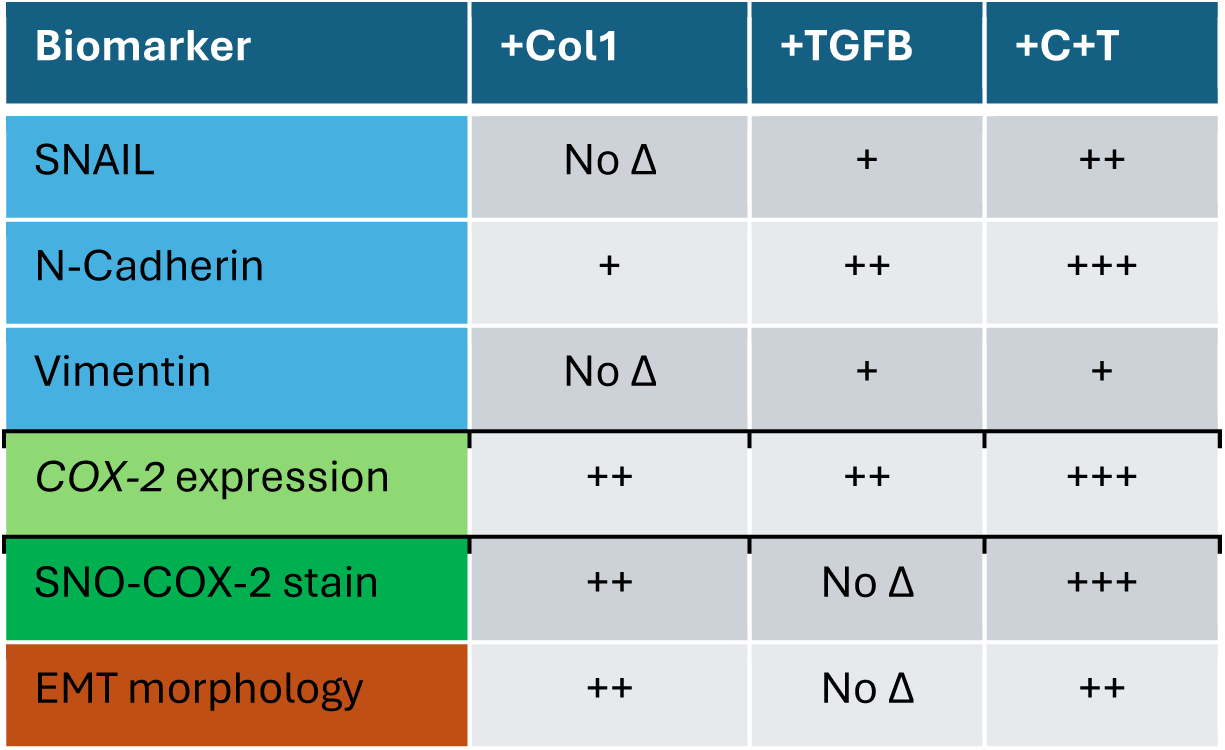
Relative changes in tumor biomarkers and tumor cell morphology. Biomarker expression is binned as no change (No Δ), small change (+), modest change (++) and large change (+++). All data are relative to the Matrigel alone conditions.

| Biomarker | +Col1 | +TGFB | +C+T |
| --- | --- | --- | --- |
| SNAIL | No $\Delta$ | + | ++ |
| N-Cadherin | + | ++ | +++ |
| Vimentin | No $\Delta$ | + | + |
| COX-2 expression | ++ | ++ | +++ |
| SNO-COX-2 stain | ++ | No $\Delta$ | +++ |
| EMT morphology | ++ | No $\Delta$ | ++ |

### SNO-COX-2 is highly expressed in mesenchymal-like regions of mouse mammary tumors

Previous work shows morphological intra-tumor heterogeneity in mouse mammary tumors derived from a D2A1 cell line isograft model, with both epithelial-like and mesenchymal-like tumor regions [30]. To assess whether SNO-COX-2 protein expression associates with a mesenchymal tumor phenotype *in vivo*, mammary tumor cells were injected into mammary fat pads of immune competent BALB/c female mice. At 3 weeks post tumor cell injection, tumors were collected for histologic and IHC analyses. In H&E-stained tumors, epithelial-like and mesenchymal-like regions were selected based on morphologic appearance only. Tumors were then IHC stained for SNO-COX-2 and vimentin, and the staining intensity was digitally quantified using stain-specific computer-assisted algorithms (**Fig 3A**). Percent SNO-COX-2 positive area was notably upregulated ∼4 fold in mesenchymal tumor regions, with a mean percent positivity of 3.42% (n=24) in epithelial regions versus 14.23% (n=26) in mesenchymal regions (**Fig 3B**, p = 0.0001). Similarly, vimentin was ∼2 fold increased in mesenchymal regions, with a mean positivity of 4.33% (n=29) for epithelial regions and 10.08% (n=32) for mesenchymal regions (**Fig. 3C**, p < 0.0001). Because COX-2 expression itself is under ovarian hormone control [31], we next controlled for potential hormonal differences between individual female mice by comparing distinct morphologic regions within the same tumor. Staining signal from epithelial and mesenchymal regions of each tumor were averaged and compared by two-way ANOVA (1-2 tumors each from six mice). Within individual tumors, there was an average ∼17-fold increase in SNO-COX-2 staining in mesenchymal tumor regions compared to epithelial tumor regions (**Fig. 3D**, p = 0.0013). Similarly, there was an average ∼2-fold increase in vimentin staining in mesenchymal regions compared to epithelial regions (**Fig. 3E**, p < 0.0001). A limitation of this mouse mammary tumor study is the lack of a validated antibody to investigate tumor levels of murine n-COX-2 in mouse tissues.

**Figure 3:**
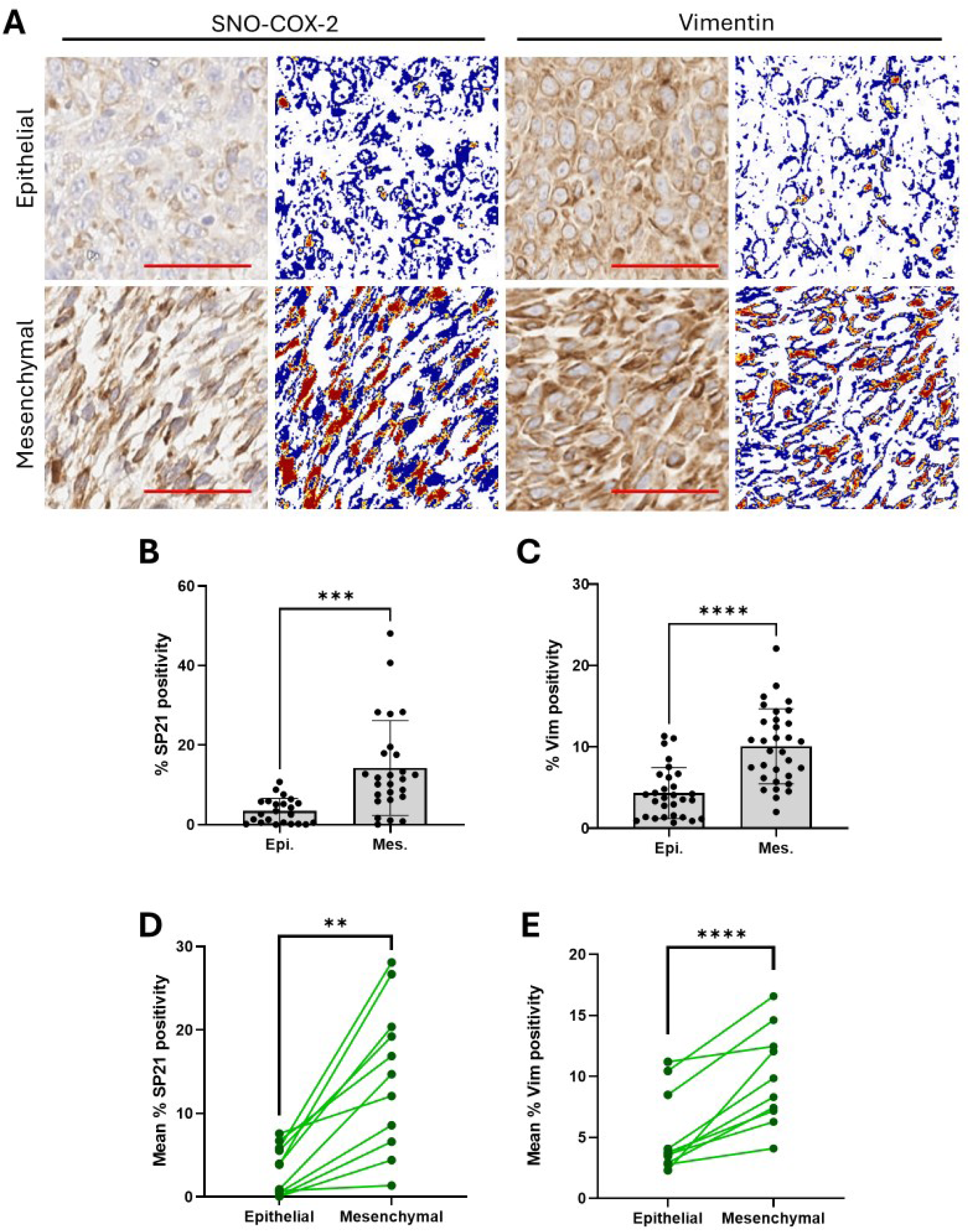
SNO-COX-2 and Vimentin staining are highest in mesenchymal regions in mouse mammary tumors with morphologic heterogeneity. **A)** Left side IHC panels show representative images of SNO-COX-2 (SP21) and vimentin staining of D2A1 isograft-induced mouse mammary tumors, delineated by epithelial and mesenchymal tumor morphologies. Positive staining is brown, nuclear counterstain is blue. Right side panels show heatmap images of SNO-COX-2 and vimentin stains derived from Aperio algorithm: background, low, medium, and high staining are indicated by blue, yellow, orange, and red, respectively. Scale bars (red) are 50 µm. **B)** Quantitation of percent positive SNO-COX-2 (% SP21) stained area from mesenchymal and epithelial regions of eleven tumors, with differences compared by unpaired t-test. **C)** As B for vimentin IHC staining across ten tumors. **D)** Intra-tumor comparison of percent positive SP21 staining area, averaged across 2-6 epithelial and 2-6 mesenchymal regions for each tumor, analyzed by two-way ANOVA. **E)** As D for vimentin IHC staining. **All)** Statistical significance is shown as *, **, *** and **** for p values < 0.05, 0.01, 0.001, and 0.0001 respectively.

Combining our *in vitro* and *in vivo* data, our results show that increased SNO-COX-2 protein expression is positively linked to the acquisition of tumor cell mesenchymal phenotype. Further, we find that both SNO-COX-2 expression and acquisition of EMT-like morphology can occur independent of EMT-related gene expression. The dominance of fibrillar type I collagen in regulating these two linked phenotypes (increased SNO-COX-2 and EMT phenotypes) implicates a key role for the tumor microenvironment in regulating SNO-COX-2. To further explore this question, we employed an omic-level, high throughput approach utilizing the microenvironment microarray (MEMA) platform [32, 33].

### MEMA assay demonstrates the importance of ECM factors in SNO-COX-2 expression and reveals combinatorial effects with soluble ligands

The MEMA platform is an omics-based, high throughput 2D cell culture assay in which cells are adhered to extracellular matrix (ECM) factors and treated with soluble ligands followed by fixation, staining, and imaging. The assay permits evaluation of effects of numerous combinations of ECM and ligands to be compared simultaneously in a high throughput *in vitro* platform. Using this platform, we evaluated 15 distinct ECM proteins paired with 20 distinct soluble ligands, resulting in 300 unique TME combinations [32, 33]. ECM factors were chosen based on published literature describing functions as either anti-or pro-tumor, with an emphasis on ECM factors related to inflammatory/regulatory microenvironments (**Table 2**). Similarly, soluble ligands were chosen based on previous studies demonstrating direct impact on COX-2 expression and/or known roles in the activation of mesenchymal cells, specifically fibroblast activation and immune cells (**Table 2**). The rationale for selecting potential mediators of mesenchymal cells is based on the fact that tumor cells can co-opt mesenchymal functions during tumor progression and thus may use these established mesenchymal pathways to upregulate tumor intrinsic SNO-COX-2.

**Table 2:** ECM factors and soluble ligands used in the MEMA assay.

| **ECM factor** | **Names** |
| --- | --- |
| COL1A | Collagen type I alpha 1 chain |
| COL3A1 | Collagen type III alpha 1 chain |
| COL5A1 | Collagen Type V Alpha 1 Chain |
| COL6A1 | Collagen Type VI Alpha 1 Chain |
| DEC | Decorin; **DCN** |
| FBN1 | Fibrillin 1 |
| FN1 | Fibronectin 1 |
| LAMA1 | Laminin Subunit Alpha 1 |
| LUM | Lumican |
| OMD | Osteomodulin |
| SPARC | Secreted Protein Acidic And Cysteine Rich; Osteonectin |
| SPP1 | Secreted phosphoprotein 1, **Osteopontin** |
| THBS1 | Thrombospondin 1 |
| TNC | Tenascin C |
| VTN | Vitronectin |
| **Ligand** | **Names** |
| 15dPGJ2 | 5-Deoxy-δ12,14-Prostaglandin J2 |
| 4HNE | 4-Hydroxynonenal |
| CCL2 | C-C Motif Chemokine Ligand 2; MCP1 |
| CTGF | Connective Tissue Growth Factor; **CCN2** |
| CXCL12 | C-X-C Motif Chemokine Ligand 12; PBSF |
| GW501516 | Cardarine |
| HGF | Hepatocyte Growth Factor |
| IFNG | Interferon Gamma |
| IL10 | Interleukin 10 |
| IL13 | Interleukin 13 |
| IL1B | Interleukin 1 Beta |
| IL4 | Interleukin 4 |
| LPS | Lipopolysaccharide; endotoxin |
| OSM | Oncostatin M |
| PBS | Phosphate Buffered Saline (control) |
| PDGFBB | Platelet Derived Growth Factor Subunit B; PDGF2 |
| PGE2 | Prostaglandin E2 |
| SAA1 | Serum Amyloid A1 |
| TGFB1 | Transforming Growth Factor Beta 1 |
| TNFA | Tumor Necrosis Factor; **TNF** |

Briefly, in triplicate 384-well plates, wells were individually coated with a single ECM factor, followed by the addition of MCF10DCIS cells. After overnight adhesion, the wells were treated with one of twenty soluble ligands, resulting in one ligand and one ECM condition per well per plate. The position of each ECM factor and ligand combination was randomized across the plate map, avoiding the outermost wells (**Sup. Fig. 3**). After 48hr of additional cell culture, cells were fixed, stained for SNO-COX-2, and cell number and SNO-COX-2 levels analyzed (**Fig. 4, Sup. Fig. 4**). We first estimated the number of cell doublings at study end, and divided cell growth into robust growth (2+ doublings over 48 hours), or reduced growth, (0-2 doublings over 48 hours or cell loss) (**Fig. 4A, Sup Fig. 5**). We observed no differences in cell numbers between plates and observed consistent SNO-COX-2 staining across our technical replicates, demonstrating technical reproducibility of the MEMA platform (**Sup. Fig. 4A-B**). The results from all 300 TME conditions (**Fig. 4B**) reveal a stark increase in SNO-COX-2 expression caused by the damage-related ECM factor Tenascin C [34] and secreted phosphoprotein 1 (SPP1; Osteopontin), regardless of the ligands paired with them. We also see high SNO-COX-2 in conditions that included the PPARdelta agonist GW501516. Interestingly, all three of these TME factors (TNC, SPP1, and GW501516) associated with significantly low cell numbers regardless of what they were paired with (**Fig. 4A, Sup. Fig. 4, Sup. Fig. 5**). In addition, several combinations with high levels of SNO-COX-2 also showed low cell numbers (**Fig. 4A-B**, black circles). Combined, these data suggest that high SNO-COX-2 protein expression associates with some aspect of growth suppression in MCF10DCIS. Of note, the PPARgamma ligand 15d-PGJ2 also induced a low cell count but had a strong negative effect on SNO-COX-2 levels. This reduction of SNO-COX-2 may not be surprising as 15d-PGJ2 is a prostaglandin known to have anti-inflammatory activity and suppress *COX-2* expression [35].

**Figure 4:**
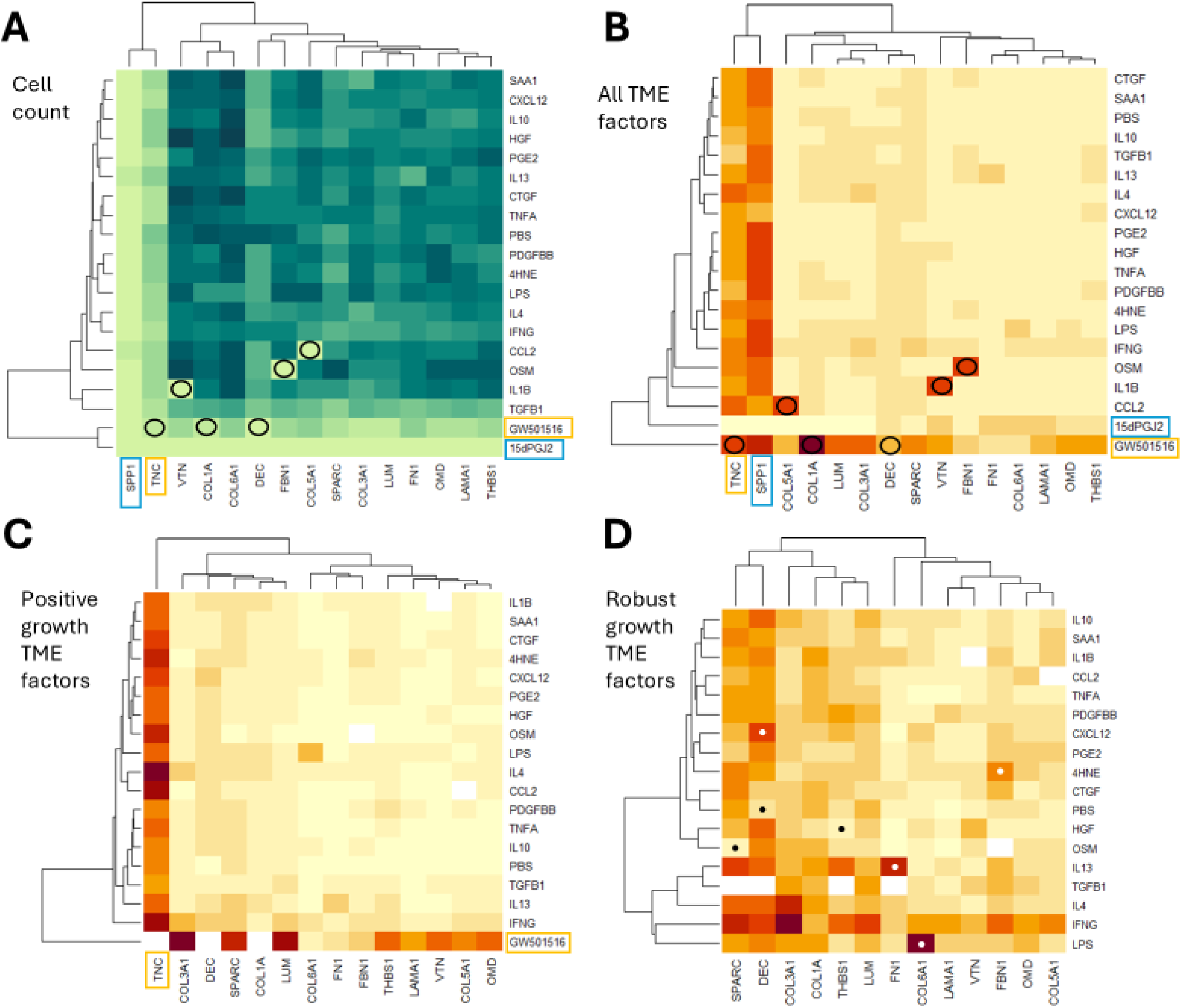
MEMA results show MCF10DCIS cell growth and SNO-COX-2 expression are under TME control. **A)** Heatmap of average cell count per image per condition in combinations of ECM factor (x axis) and soluble ligand (y axis); darker colors indicate higher cell count. Briefly, each well was imaged nine times and the cell count across all three technical replicates was averaged to a single value. All SPP1as well as 15dPGJ2 conditions had negative growth (<0 doubling) and are highlighted with (blue boxes) and unique combinations with negative growth are depicted with black circles. Similarly, all TNC and GW501516 associated with low growth (0-2 doublings) and are indicated by orange boxes. **B)** As A, but showing average SP21 (SNO-COX-2) staining per well per condition. **C)** As B, excluding conditions with negative growth. **D**) As C, excluding conditions with low growth (<2 doublings). White dots and black dots highlight examples of combinations of TME factors that induce high or low SNO-COX-2 expression (respectively) compared to the rest of the row/column.

In the TME conditions with high cell numbers (robust cell growth) (**Fig. 4D**), ECM factors were a stronger influence on the nitrosylation of COX-2 (SNO-COX-2) than soluble ligands. SPARC, Decorin, Type I and III fibrillar collagens, Thrombospondin 1 and Lumican significantly elevated SNO-COX-2 protein levels. By contrast, SNO-COX-2 levels were low in wells coated with type V and VI collagens, laminin 1 and vitronectin. Among the soluble ligands, interferon gamma (INFG; IFN-γ) stands out as a strong inducer of SNO-COX-2. With respect to SNO-COX-2 staining, INFG clustered with other immune regulators, IL-13, IL-4, and TGFβ-1, as well as LPS (endotoxin), all of which showed increased SNO-COX-2 protein levels in multiple conditions (**Fig. 4D**). Every factor had at least one partner that caused drastically higher or lower SNO-COX-2 protein compared to their other pairings. Examples of high SNO-COX-2 conditions include COL6A1+LPS, FN1+IL-13, FBN1+4HNE, and DEC+CXCL12 (**Fig. 4D**, white dots). Examples of low SNO-COX-2 conditions include THBS1+HGF, SPARC+OSM, and DEC+PBS (**Fig. 4D**, black dots). These results highlight the importance not just of individual TME factors but of factor combinations in the control of COX-2 nitrosylation.

In summary, our MEMA model has revealed three cellular phenotypes that associate with altered SNO-COX-2 levels. First, a phenotype of reduced cell growth or cell loss with highest SNO-COX-2 expression, notably induced by two damage-associated ECM proteins and a PPARdelta agonist. Second, a phenotype of robust cell growth in which EMT factors associated with fibrillar collagens and ligands associated with immune induction and resolution that also promote COX-2 nitrosylation. Third, a phenotype of robust growth but suppressed SNO-COX-2 expression associated with factors which may represent a tumor-suppressive or normalizing microenvironment.

## Discussion

COX-2 is an essential inflammatory regulator that is also compellingly involved in tumor biology, including tumor cell invasion and EMT [3–7, 9, 10, 18, 22]. However, tumor COX-2 biology is insufficiently understood, impacting its usefulness as a target for prognostics, prevention or treatment. Here we address a specific post-translational modification of the COX-2 protein, S-nitrosylation of Cys-526, to ask if tumor cell expression of the nitrosylated form associates more strongly with tumor cell invasion and/or EMT-like phenotypes compared to the non-nitrosylated form. We found SNO-COX-2 protein levels, but not non-nitrosylated COX-2 levels, correlated strongly with EMT-like invasive morphology in *in vitro* and *in vivo* breast cancer models. SNO-COX-2 protein levels also associated more strongly with invasive morphology than did gene expression of the EMT biomarkers *SNAIL*, vimentin and N-cadherin. Further, we identified strong microenvironmental control of SNO-COX-2 protein expression, specifically by TME factors associated with wound healing and tumor microenvironments. In sum, SNO-COX-2 is a nitrosylated form of COX-2 that is not widely recognized nor examined in the COX-2 literature but holds potential as a breast cancer biomarker and therapeutic target.

In a 3D tumor spheroid model, SNO-COX-2 protein expression associated closely with invasive morphology of MDF10DCIS cells, a cell line that models early-stage human breast cancer. We expected classical markers of EMT to also tract with invasive morphology. Instead, we found a dissociation between gene expression of EMT biomarkers *SNAIL*, vimentin, N-cadherin, and *COX-2* and mesenchymal morphology, with only SNO-COX-2 protein levels having a strong correlation with mesenchymal phenotypes. Specifically, invasive MCF10DCIS morphology, driven by type I collagen, occurred in the absence of *SNAIL,* vimentin, and N-cadherin gene induction, whereas TGFβ-1 strongly induced these EMT genes despite the absence of invasive phenotypes. These data may suggest forms of EMT distinct from dependence on classic transcription factors, and/or add to the fields of partial EMT and epithelial/mesenchymal plasticity (EMP) [24]. It has previously been reported that cells in partial EMT are critical for metastasis in a mouse breast cancer model, and that partial EMT cells vacillate between epithelial and hybrid EMT [36]. Our data suggest a specific role for SNO-COX-2 in partial EMT, as defined by mesenchymal morphology in the absence of EMT biomarker transcription.

Our microenvironment microarray cell culture platform allowed for the investigation of 300 combinations of ECM proteins and soluble immune-regulatory ligands, which revealed complex microenvironmental control of SNO-COX-2 protein levels. Using this MEMA platform, we find that ECM proteins classically associated with early wound healing were dominant, having stronger induction effects on SNO-COX-2 expression than soluble inflammatory ligands. For example, ECM proteins known to be tissue damage signals [34], inducing inflammation through toll-like receptor signaling and the NF-kB signaling pathway, strongly induced SNO-COX-2. These ECM proteins included decorin, fibrinogen, tenascin C [34], and SPARC [37]. SNO-COX-2 was also highly upregulated by the early inflammatory ligand interferon gamma, which can induce NF-kB signaling [38]. Surprisingly, several additional inflammatory ligands that are under NF-kB control, including prostaglandin PGE_2_, IL-1B, TNF-a, and CCL2 did not upregulate SNO-COX-2 in our model, highlighting the complexity of SNO-COX-2 expression in this single cell line. The MEMA data also revealed that SNO-COX-2 is highly upregulated by early ECM damage signals and interferon gamma, as well as ECM (fibrillar type I and III collagens) and immune-regulatory ligands associated with the recruitment of fibroblasts that occurs following the initial inflammatory damage response. Whether SNO-COX-2 plays a key role in early wound/damage response remains an open question.

As wound healing progresses, inflammation turns from pro-inflammatory to regulatory due to upregulation of immune suppressive/regulatory cytokines. In turn, chronic immune suppression and avoidance is a hallmark of cancer [1, 2]. We find that SNO-COX-2 was also upregulated by regulatory cytokines (IL-4 and IL-13 [39]), especially in conjunction with early wound-healing ECM proteins described above. Further, ECM components reported to regulate and resolve inflammation, including thrombospondin 1 and lumican, also induced SNO-COX-2. Combined, we find immune regulatory signals classically associated with wound resolution and tumor cell immune avoidance upregulate SNO-COX-2 in our breast cancer model. These data raise the intriguing but untested possibility that tumor-cell intrinsic SNO-COX-2 may play a role in immune avoidance.

Of note and contrary to our 3D culture findings, the regulatory cytokine TGFβ-1 [40] also induced high SNO-COX-2 expression in the (2D) MEMA assay, clustering with regulatory cytokines IL-4 and IL-13. This suggests distinct effects of TGFβ-1 on MCF10DCIS cells in 2D versus 3D cell culture conditions, which may be attributed to laminin. In the MEMA assay, we identified laminin 1, a major non-collagen component of basement membranes, as highly suppressive of SNO-COX-2. Laminin 1 is a major component of Matrigel and is present at high concentration in the 3D but not the 2D culture models. Further, laminin 1 can suppress tumor cell invasion in 3D culture [41]. Since the expression of SNO-COX-2 induced by TGFβ-1 in the MEMA was lowest when TGFβ-1 was paired with laminin 1, the suppressive effect of laminin 1 may be dominant over TGFβ-1 when TGFβ-1 is combined with laminin. ECM suppression of SNO-COX-2 was not limited to laminin. Another ECM factor that reduced SNO-COX-2 expression was collagen VI, a non-fibrillar collagen associated with tissue elasticity [42]. Suppression of SNO-COX-2 by collagen VI is consistent with earlier studies demonstrating that decreased collagen tension associates with epithelial organization of tumor cells (e.g. mesenchymal-to-epithelial transition; MET) while high collagen tension, such as provided by fibrillar type I and III collagens, promotes EMT [23, 43].

The MEMA assay revealed another aspect of SNO-COX-2 regulation in MCF10DCIS cells, which is that high SNO-COX-2 was strongly induced by ECM and ligands that also suppressed cell growth. Low cell counts in the 2D MEMA assay could be caused by an increase in cell death, raising the possibility that SNO-COX-2 has a role in cell death, however, we did not observe correlations between high SNO-COX-2 and cell death in our 3D assay (**Sup. Fig. 1E-F**). Alternatively, because EMT is frequently associated with reduced proliferation, SNO-COX-2 may play a role in the proliferation versus motility dichotomy of EMT. Three factors associated with low cell counts that induced dramatically high expression of SNO-COX-2 were Tenascin C, SPP1 (Osteopontin), and GW501516 (cardarine). Tenascin C and SPP1 are known ECM components capable of inducing EMT-like changes in tumor cells [44, 45], while GW501516 is a PPARdelta agonist that enhanced EMT in melanoma cells [46]. Because the three factors associated with high SNO-COX-2 and low cell count are all related to EMT, the observed low cell count may be EMT-linked. Potential roles for SNO-COX-2 in mediating the proliferation/motility dichotomy remain interesting targets of further study. Finally, the TME conditions that led to high SNO-COX-2, EMT morphology, and low cell count in the 3D assays had increased cellular debris. Further research is needed to understand the nature of this cell debris, which could be remnants of cell death and/or extracellular vesicles.

Strengths of our study include the focus on an understudied form of COX-2, SNO-COX-2, in breast cancer mesenchymal phenotypes and EMT; our use of a relevant early-stage human breast cancer cell line model that readily permits TME control of tumor cell invasive morphology; the breadth of microenvironmental factors that we tested for their ability to regulate SNO-COX-2 protein levels; and the employment of validated antibodies for n-COX-2 and SNO-COX-2. An additional strength of our work is the relevance of our findings to the field of tumor-associated COX-2 research in general. Our study also has weaknesses, including the singular cell line used, a selection driven by the unique DCIS attributes of this cell line. While it is possible that our findings are specific to MCF10DCIS, we addressed this limitation in part with our examination of murine mammary D2A1 tumor cell isografts. In future work, additional cell lines and cancer cell types should be examined. Colon cancer lines would be important, based on the body of literature showing COX-2 as an oncogene and target of treatment [6, 47]. Second, our MEMA study is exploratory, with a single experiment performed in technical triplicate. Further, we cannot disregard potential concerns regarding the quality of the thirty-five different commercially available reagents utilized (**Table 2**), the quality of which could influence some MEMA results. We note as a potential example that fibronectin (FN1), despite being a pro-tumor ECM protein associated with metastasis, induced the lowest expression of SNO-COX-2 of any ECM protein. While it is possible that fibronectin downregulates SNO-COX-2 in the presence of most ligands, it may also be that our plated fibronectin was in some way defective. Also, the MEMA platform is a 2D model, which reduces its biological relevance. To address these weaknesses, the MEMA study could be repeated and the combinations of factors that show promise explored in 3D culture assays. Finally, a study weakness is the descriptive nature of the data given that we did not determine a causal relationship between SNO-COX-2 and tumor cell mesenchymal/invasive phenotypes. To definitively test whether SNO-COX-2 is necessary to induce invasive morphology, a mutant form that is unable to be S-nitrosylated on Cys-526 will be required.

In conclusion, our observations support the utility of SNO-COX-2 as a tumor invasion biomarker upregulated in microenvironmental conditions that are classically associated with early wound responses, wound resolution, and pro-tumor, while being suppressed by environments that are associated with anti-tumor, tissue normalization. Thus, tumor-intrinsic SNO-COX-2 may be an untapped biomarker for more invasive, EMT-like cancer cell phenotypes that may prove to be a superior prognostic marker when compared to non-nitrosylated COX-2 or classical biomarkers of EMT, topics that deserve further investigation. Additionally, while SNO-COX-2 appeared independent of iNOS in our cell culture model, COX-2 and iNOS are both biomarkers of poor prognosis in ER-breast cancer [48, 49]. COX-2 and iNOS have been shown to be capable of upregulating each other’s expression, and inhibition of iNOS and its homologues is being explored as a therapeutic treatment in breast cancer [14, 50, 51]. It is intriguing to speculate that efficacy of iNOS inhibitors may be causally linked to suppression of SNO-COX-2, suggesting that future detection of SNO-COX-2 may inform use of NSAIDS and nitric oxide pathway inhibitors in cancer treatment. In addition, our results suggest that distinguishing between SNO-COX-2 and n-COX-2 may substantially improve our understanding of COX-2 in general, as recently noted by researchers in the field [52, 53].

## Supporting information

Full Supplemental File

## Glossary

1400W: N-[[3-(aminomethyl)phenyl]methyl]-ethanimidamide, dihydrochloride, a selective inhibitor of iNOS (NOS2).
AML2: Acute Myeloid Leukemia 2, AKA **OCI-AML-2**; an acute myeloid leukemia tumor cell line.
*CDH2*: Cadherin-2 gene, AKA *NCAD* or N-cadherin. Encodes a transmembrane protein (N-cad; N-cadherin) which forms adherens junctions in mesenchymal cells. Common EMT marker.
+Col1, +C: 3D culture conditions including fibrillar type I collagen.
COX-2: Cyclooxygenase-2; the gene (*COX-2*) or the protein (COX-2). AKA prostaglandin-endoperoxide synthase 2 (*PTGS2*).
CX229: Antibody clone used in histology which detects human COX-2 that is not S-nitrosylated on Cys-526.
+C+T: +Col1+TGFB; 3D culture conditions including both fibrillar type I collagen and soluble TGFβ-1.
D2A1: Mouse mammary cancer cell line derived from spontaneous mammary tumors.
DAPI: 4’,6-diamidino-2-phenylindole, a fluorescent dye that binds DNA for nuclear imaging.
DCIS: (Mammary) Ductal Carcinoma *In Situ* – not to be mistaken for the cell line MCF10DCIS.
ECM: Extracellular Matrix.
EMP: Epithelial/Mesenchymal Plasticity, an alternative name for partial EMT. Can also include MET.
EMT: Epithelial-to-Mesenchymal Transition.
eNOS: Endothelial Nitric Oxide Synthase, AKA **NOS3**.
FFPE: Formalin-fixed paraffin-embedded; common tissue preservation approach for histology.
H&E staining: Hematoxylin and Eosin staining.
IDC: (Mammary) Invasive Ductal Carcinoma.
iNOS: Inducible Nitric Oxide Synthase, AKA **NOS2**.
L-NMMA: N^G^-Methyl-L-arginine, a relatively non-selective nitric oxide synthase (NOS) inhibitor.
MCF10DCIS: An oncogene-transformed spontaneously immortalized human breast cell line capable of forming DCIS-like lesions when injected into the fat pad of mice. AKA: MCF10ADCIS.com, MCF10DCIS.com, DCIS.com, etc.
MEMA: Microenvironment Microarray.
MET: Mesenchymal-to-Epithelial Transition.
*NCAD,* N-cad: See *CDH2*.
nNOS: Neuronal Nitric Oxide Synthase, AKA **NOS1**.
NSAID: Non-Steroidal Anti-Inflammatory Drug (e.g. aspirin, ibuprofen).
n-COX-2: COX-2 which has <u>not</u> been S-nitrosylated on Cys-526.
Mgel: Matrigel, trade name (Corning Life Sciences); extracellular matrix material derived from Engelbreth-Holm-Swarm mouse sarcoma, enriched in laminin 1 basement membrane protein.
PDAC: Pancreatic Ductal Adenocarcinoma.
PGH_2_: Prostaglandin H_2_, the product of COX-2 and a precursor to other prostaglandins and related lipids.
*SNAIL*: AKA *SNAI1*, gene for SNAIL, a major upstream mesenchymal transcription factor commonly used as a biomarker of EMT.
SNO-COX-2: COX-2 which has been S-nitrosylated on Cys-526.
SP21: Antibody clone used in histology which detects SNO-COX-2.
+TGFB, +T: 3D culture conditions including soluble TGFβ-1.
*TGFB1*, TGFβ-1: Transforming growth factor beta 1; the gene (*TGFB1*) or the protein (TGFβ-1).
TME: Tumor Microenvironment.
*VIM*: Vimentin gene, encodes a structural an intermediate filament expressed in mesenchymal cells. Common EMT marker.

## Acknowledgements

This work was supported by funds from the Cancer Early Detection Advanced Research (CEDAR) Center (ID # 6370919) and the Knight Cancer Institute CCSG (NIH P30 CA069533) to PS and JK, NIH R01CA169175 to PS, Breast Cancer Research Foundation (BCRF) to PS, Willard L. and Ruth P. Eccles and Leonard Schnitzer Family Foundations to PS, NIH T32 CA254888 to RJH, and Kuni Foundation to RJH and PS. MCF10DCIS cells were a gift from K. Polyak (Harvard School of Medicine, Boston, MA). AML2 cells were a gift from Anupriya Agarwal (OHSU). We thank Dr. Nathan Pennock for assistance with the MEMA study design, Dr. Michelle Ozaki, Elise DeWilde, Moqing Liu, Katlyn Devlin, and Cydney Hunt for technical assistance, Dr. Sudarshan Anand for qPCR advice and reagents, and Solange Bassale and the Knight Biostatistics Shared Resource (NIH P30 CA069533) for statistics support. The research reported in this publication used computational infrastructure supported by NIH Award Number S10OD034224. We acknowledge support and expert technical assistance by staff in the Advanced Multiscale Microscopy Shared Resource (NIH P30 CA069533) and the OHSU University Shared Resources Program. It takes a team.

## Methods

### Cell lines & cell culture, general

MCF10DCIS (aka: MCF10ADCIS.com) cells were a gift from K. Polyak (Harvard School of Medicine, Boston, MA). The D2A1 cell line was a gift from Ann Chambers (Ontario, Canada). A cell pellet of AML2 (OCI-AML-2) for RNA isolation was generously provided by Anupriya Agarwal (OHSU, Portland, OR). Cells were grown in DMEM/F12 (1:1) medium supplemented with L-glutamine and HEPES (HyClone SH30023.01) with the following additions: Horse Serum, non-heat-inactivated (Hyclone SH30074.03), to a final concentration of 5%; Insulin (Gibco 12585-014) to a final concentration of 10 ug/mL; Hydrocortisone (Sigma H-4001) to a final concentration of 500 ng/mL; recombinant human EGF (BD354052) to a final concentration of 20 ng/mL; Cholera Toxin (Sigma C9903) to a final concentration of 100 ng/mL, all as previously described [54]. In 2D culture, cells were passaged on cell-treated culture plates, at 37°C with 5% CO, fed every two days and passaged when cells were at 60-80% confluency, using trypsin.

### Cell culture, 3D

Briefly, for 3D culture experiments, MCF10DCIS cells were suspended in Matrigel (Corning 354234) Matrigel lots used in these experiments were 0323002, 1035003, or 26323002. All technical replicates of a given experiment used the same lot. Matrigel was thawed on ice and diluted to 8 mg/mL with standard growth media. For collagen-containing wells, rat tail collagen (Corning 354249) was added to a neutralization solution of 10x PBS, 1N NaOH, and growth media to a final collagen concentration of 4 mg/mL, a final PBS concentration of 1x, and a final equivalent NaOH concentration of 0.0075 N, adjusted to a final pH of 7.0 – 7.5 with HCl if needed. Prior to the addition of collagen to the diluted Matrigel, the collagen solution was incubated on ice for 1 hour to allow neutralization, which is required for fibril formation. On ice, de-adhered cells were resuspended to a final concentration of 60k cells/mL with either Matrigel alone (8 mg/mL) or Matrigel and collagen at a 30:70 ratio by volume (final concentrations of 2.4 mg/mL matrigel and 2.8 mg/mL collagen). 250 µL of the culture mixtures were plated per well onto the filters of sterile 0.4 µm pore transwell inserts (Corning 353095) in sterile 24-well cell culture plates (Corning 3526). The resulting gels containing cells were incubated (37°C, 5% CO_2_) for 45 minutes to allow solidification, after which 250 µL of growth media was carefully added to the transwell on top of the filter and 700 µL of growth media was added to each well under the filter. Empty wells were filled with sterile PBS to prevent evaporation. Twenty-four hours after initial plating (Day 1), and every forty-eight hours after (Day 3, 5, etc.), cultures were fed by removing 125 µL (roughly half) of the media from inside the transwell and replacing it with 125 µL of fresh media. For cultures treated with TGFβ-1, the replacement media also included 5 ng/mL TGFβ-1, for a final concentration of 2.5 ng/mL in the transwell. Media under the transwell inserts was not changed. For histology, wells were carefully emptied of media both inside and out of the transwell, gently washed twice with PBS, then refilled with 5% paraformaldehyde (PFA) in PBS, after which plates were sealed with parafilm and incubated at RT for 20 minutes. Fixed gel pads were rinsed twice with PBS and kept in 70% ethanol (at RT, sealed and in the dark) until embedded; at least overnight. For embedding, pads were gently removed from transwells by cutting out the filter with a razor blade and placing into a histo-cassette prior to paraffin embedding. For imaging, paraffin-embedded samples were sectioned into 4-micron sections and mounted on glass microscopy slides.

### Mouse tumor model

Mouse mammary tumors were derived from the murine mammary tumor cell line D2A1, orthotopically injected into mammary glands of immunocompetent Balb/c mice, as previously described [18]. Briefly, 2 x 10^4^ D2A1 cells per gland were injected into the fourth mammary glands of BALB/c females (12-13 weeks of age) and after three weeks of growth, tumor-bearing mammary glands were collected, formalin fixed, paraffin embedded, and thin sectioned for histologic-based assays. Mammary tumors from six mice were sectioned and used for SNO-COX-2 and vimentin IHC staining.

### Histology

All histology was performed using slide-mounted 4 µm tissue sections. Briefly, slides were de-paraffinized and washed in xylene, then sequentially re-hydrated with ethanol gradient

concentrations. For immunofluorescent (IF) staining of 3D cell cultures, thin sectioned slides were treated with antigen retrieval solution (Dako S1699) in a pressure cooker (Dako Pascal S2800) at 125°C for 5 min, followed by treatment with blocking buffer (Biocare BS966L) for 1 hour at room temperature. Primary antibodies for anti-SNO-COX-2, clone SP21 (1:50, Thermo Fisher scientific RM-9121) and anti-n-COX-2, clone CX229 (1:100, Cayman Chemicals 160112), were incubated overnight on each sample, followed by wash steps. Secondary antibodies were applied: donkey anti-rabbit 555 (1:250, Thermofisher A-31572?) and goat anti mouse 647 (1:250, Thermofisher A-21236) for 2 hours at room temperature, protected from light, washed, followed by addition of DAPI (1:50, 5mg/ml) for 10 min at room temperature.

For Immunohistochemical (IHC) assessment of the D2A1 mouse mammry tumors, slides with tumors from six mice were blocked in 3% hydrogen peroxide in methanol for 15 minutes, washed twice with TBST, and blocked for an additional 15 minutes with Background Sniper (Biocare Medical, BS966) followed by staining using anti-SNO-COX-2 primary clone SP21 (1:50, Thermo Fisher scientific RM-9121) at 4°C overnight or anti-vimentin primary (1:400, 5741S, Cell Signaling Technology) at RT for 2 hours. After washes, Histofine anti-rabbit secondary for mouse (Nichirei 414341F) was applied for 30 minutes at RT. After washes, slides were developed using by 3,3′-diaminobenzidine (DAB; Novus Biologicals, #SK-4105-NB). Vimentin IHC staining was performed as per the last paragraph using 1:1000 primary antibody concentration. Counterstaining was performed with hematoxylin. Slides were then dehydrated, coverslipped, and imaged.

### Imaging and analysis

For histologic assessment of 3D spheroid tumor phenotypes, H&E images were binned by similar morphological types and number of each morphology “events” captured by manual counting by two blinded and independent reviewers. The morphologic categories were: 1) Smooth edged spherical which had no unusual protrusions or invasive morphology; 2) Partially invasive spheroids which were largely round but possessed at least one cellular extension or protrusion; 3) Multicellular invasive spheroids which lacked spherical shape and possessed multiple cellular extensions or protrusions resulting in spindle-like or star-like shapes.; and 4) Single-invasive cells were single cells with spindle-like shape or many protrusions, consistent with EMT. Scored events were pooled across replicate by experiment and compared as percentage values of experimental events and analyzed via mixed-effects modeling.

For If studies, IF stained images were captured using an Zen Observer.Z1 microscope with ApoTome.2 attachment. Stain signals were analyzed using Zeiss’s Zen software and average stain intensity by area recorded. Average intensity by area was normalized against the average intensity across all matrigel-alone condition events for a given imaging experiment. Normalized events were averaged within each technical triplicate. Results were compared by two-way ANOVA (experiment vs condition) followed by Tukey’s multiple comparisons test between each different condition (alpha = 0.05). IHC and H&E images were taken using an Aperio ScanScope AT (Leica Biosystems) and image analysis was carried out using Aperio ImageScope software. For mouse tumor IHC images stained for vimentin and SNO-COX-2, 1-6 tumor regions (regions of interest, ROI) per tumor were chosen based on histologic mesenchymal- and epithelial-like characters. Color-temperature algorithms were designed for each stain to exclude background and the % positive stained area divided by the total ROI size to obtain percent area stained. ROIs were pooled by morphologic type, i.e. epithelial or mesenchymal, and analyzed by unpaired t-test. For a subset analysis of intra-tumor comparisons, ROIs were grouped by tumor and by morphologic subtype and analyzed by two-way ANOVA (α = 0.05).

### RNA isolation and RT-qPCR

After 5 days in culture, spheroids in matrigel pads (as well as pelleted AML2 cells) were digested by TRIzol LS reagent (Invitrogen 10296028). Briefly, media was removed and 750 mL TRIzol LS added per well, followed by trituration (repeatedly pipetting up and down) of each pad until dissolved. After 3 min RT incubation and centrifugation to remove excess matrix components, resulting lysate was spun through QiaShredder columns (Qiagen 79656) followed by centrifugation at 12,000 g for 10 min at 4°C to pellet excess matrix material. To the resulting supernatant, 200 µL of chloroform was added, followed by 5-10 forceful tube inversions, 15 seconds of vortexing, a 3 min RT incubation, and centrifugation at 12,000 g for 15 min at 4°C to separate phenol and chloroform phases. The RNA-containing (upper) phase was collected and mixed with an equal volume of 100% ethanol, which was used as input for purification by a Direct-zol RNA MicroPrep kit (Zymo Research R2060). Eluted RNA was converted into cDNA 1:1 using SuperScript IV VILO MasterMix (Invitogen #11756050). 1 ng of cDNA was used in each RT-qPCR replicate. RT-qPCR was performed using SSoAdvanced Universal SYBR Green Supermix (Bio-Rad 1725271) in 384-well PCR plates (Bio-Rad #MSP3842) sealed with MicroAmp Optical Adhesive Film (Applied Biosystems 4311971) using a CFX Opus 384 Real-Time PCR System (Bio-Rad 12011452) and recommended settings. Primers for RT-qPCR were designed according to SuperScript kit instructions and validated against purified RNA from 2D cell culture pellets (MCF10ADCIS or AML2). The list of primers in provided in **Sup. Table 3**. For negative controls, we utilized no-template and no-reverse-transcriptase “cDNA” samples.

Reported results are from three 3D culture experiments. Each 3D culture experiment included three technical replicates (wells; gel pads) of each condition from which RNA was prepared; RNA was not pooled between culture technical replicates. Each RT-qPCR reaction was run once using RNA from each technical replicate in technical triplicate (three qPCR wells per culture well/pad); ΔCq values were calculated as the difference between average Cq values of a given primer and the GAPDH housekeeper. ΔΔCq values were calculated as the difference between average ΔCq values for a given primer across the three culture replicates of a given experimental condition and the average ΔCq value of the matrigel-alone culture replicates. Experiments were compared using 2-way ANVOA (condition vs. experiment) followed by Tukey’s multiple comparisons test between each different condition (α = 0.05).

### Microenvironment Microarray (MEMA)

MEMAs were performed as described previously [32, 33]. ECM and ligand factors can be found in **Sup. Table 2**. Briefly, the plate layout for 384-well MEMA plates (**Sup. Fig. 3A**) excluded the outer wells (i.e. rows A and P, columns 1 and 24) to avoid edge effects. Wells A2 through A23 were used as plating/staining cell-line postive controls, which received COL1A+PBS. All other outer wells and the inter-well spaces were filled with sterile PBS to prevent evaporation. The remaining 308 inner wells (B-O, 2-23) each featured a single ECM/ligand combination, plus five additional wells of COL1A+PBS controls and two wells lacking ECM and ligand as additional plating controls. Three plates with identical layouts were run, resulting in three technical replicates for each experiment condition.

ECM factors were plated into the bottom of the wells of 384-well plates (ThermoFisher 242764) according to the plate map (**Sup. Fig. 3**, top) at a concentration per surface area of 1 µg/cm^2^, followed by plate centrifuged (10 sec, 1000 rpm), and room temperature drying time in sterile conditions for two days. MCF10DCIS cells suspended in normal growth media (see cell culture) and plated at ∼1050 cells in 100 µL per well (10.5k cells/mL). HCA-7 colon cancer cells, as a positive SNO-COX-2 cell line [16], were plated in line control wells (A2-A23) in normal growth media (see cell culture) at ∼2000 cells in 100 µL per well (20k cells/mL). Plates were incubated overnight (37°C, 5% CO_2_) to allow cell adherence, then 100 µL of media was added supplemented with PBS (control) or a soluble ligand according to the plate map (**Sup. Fig. 3**, bottom) to a final volume of 200 µL and final concentrations listed in **Sup. Table 2**. Plates were incubated for 48 hours, then fixed with 4% PFA and washed with PBS three times. Adhered cells were stained for SNO-COX-2 as described in IF staining (see histology) and with DAPI for nuclear visualization and counted.

Imaging of MEMA plates was performed using a laser-scanning confocal microscope LSM880 equipped with AiryScan. A 3×3 grid of non-overlapping representative images were taken from each well. Cells were segmented and fluorescent intensity calculated for each stain as previously reported [32], using CellPose V1 in lieu of Cell Profiler. Cell counts were averaged across all images of all three replicate plates. Cell doublings were estimated based on initial plating density. Average SNO-COX-2 intensity per cell per image were scored and the median image per well (excluding images without cells) was chosen as representative for the well for staining intensity. Stain medians per well were averaged together across replicates to generate the experimental mean.

## Citations

1. Coussens, L.M. and Z. Werb, Inflammation and cancer. Nature, 2002. 420(6917): p. 860–7.

2. Hanahan, D. and R.A. Weinberg, Hallmarks of cancer: the next generation. Cell, 2011. 144(5): p. 646–74.

3. Nakanishi, M. and D.W. Rosenberg, Multifaceted roles of PGE2 in inflammation and cancer. Semin Immunopathol, 2013. 35(2): p. 123–37.

4. Ristimaki, A., et al., Prognostic significance of elevated cyclooxygenase-2 expression in breast cancer. Cancer Res, 2002. 62(3): p. 632–5.

5. Richardsen, E., et al., COX-2 is overexpressed in primary prostate cancer with metastatic potential and may predict survival. A comparison study between COX-2, TGF-beta, IL-10 and Ki67. Cancer Epidemiol, 2010. 34(3): p. 316–22.

6. Gupta, R.A. and R.N. Dubois, Colorectal cancer prevention and treatment by inhibition of cyclooxygenase-2. Nat Rev Cancer, 2001. 1(1): p. 11–21.

7. Liu, C.H., et al., Overexpression of cyclooxygenase-2 is sufficient to induce tumorigenesis in transgenic mice. J Biol Chem, 2001. 276(21): p. 18563–9.

8. Rao, P. and E.E. Knaus, Evolution of nonsteroidal anti-inflammatory drugs (NSAIDs): cyclooxygenase (COX) inhibition and beyond. J Pharm Pharm Sci, 2008. 11(2): p. 81s–110s.

9. Jacoby, R.F., et al., The cyclooxygenase-2 inhibitor celecoxib is a potent preventive and therapeutic agent in the min mouse model of adenomatous polyposis. Cancer Res, 2000. 60(18): p. 5040–4.

10. Arun, B. and P. Goss, The role of COX-2 inhibition in breast cancer treatment and prevention. Semin Oncol, 2004. 31(2 Suppl 7): p. 22-9.

11. Peng, C., et al., Regular aspirin use, breast tumor characteristics and long-term breast cancer survival. NPJ Breast Cancer, 2025. 11(1): p. 62.

12. Alexanian, A. and A. Sorokin, Cyclooxygenase 2: protein-protein interactions and posttranslational modifications. Physiol Genomics, 2017. 49(11): p. 667–681.

13. Kim, S.F., D.A. Huri, and S.H. Snyder, Inducible nitric oxide synthase binds, S-nitrosylates, and activates cyclooxygenase-2. Science, 2005. 310(5756): p. 1966–70.

14. Granados-Principal, S., et al., Inhibition of iNOS as a novel effective targeted therapy against triple-negative breast cancer. Breast Cancer Res, 2015. 17(1): p. 25.

15. Glynn, S.A., et al., Increased NOS2 predicts poor survival in estrogen receptor-negative breast cancer patients. J Clin Invest, 2010. 120(11): p. 3843–54.

16. Jindal, S., et al., S-nitrosylated and non-nitrosylated COX2 have differential expression and distinct subcellular localization in normal and breast cancer tissue. NPJ Breast Cancer, 2020. 6(1): p. 62.

17. Jindal, S., et al., Characterization of weaning-induced breast involution in women: implications for young women’s breast cancer. NPJ Breast Cancer, 2020. 6: p. 55.

18. Lyons, T.R., et al., Cyclooxygenase-2-dependent lymphangiogenesis promotes nodal metastasis of postpartum breast cancer. J Clin Invest, 2014. 124(9): p. 3901–12.

19. Miller, F.R., et al., MCF10DCIS.com xenograft model of human comedo ductal carcinoma in situ. J Natl Cancer Inst, 2000. 92(14): p. 1185–6.

20. Hu, M., et al., Regulation of in situ to invasive breast carcinoma transition. Cancer Cell, 2008. 13(5): p. 394–406.

21. Russell, T.D., et al., Myoepithelial cell differentiation markers in ductal carcinoma in situ progression. Am J Pathol, 2015. 185(11): p. 3076–89.

22. Lyons, T.R., et al., Postpartum mammary gland involution drives progression of ductal carcinoma in situ through collagen and COX-2. Nat Med, 2011. 17(9): p. 1109–15.

23. Maller, O., et al., Collagen architecture in pregnancy-induced protection from breast cancer. J Cell Sci, 2013. 126(Pt 18): p. 4108–10.

24. Yang, J., et al., Guidelines and definitions for research on epithelial-mesenchymal transition. Nat Rev Mol Cell Biol, 2020. 21(6): p. 341–352.

25. Shintani, Y., et al., Collagen I promotes epithelial-to-mesenchymal transition in lung cancer cells via transforming growth factor-beta signaling. Am J Respir Cell Mol Biol, 2008. 38(1): p. 95–104.

26. Xu, J., S. Lamouille, and R. Derynck, TGF-beta-induced epithelial to mesenchymal transition. Cell Res, 2009. 19(2): p. 156–72.

27. Derynck, R., R.J. Akhurst, and A. Balmain, TGF-beta signaling in tumor suppression and cancer progression. Nat Genet, 2001. 29(2): p. 117–29.

28. Debnath, J., et al., The role of apoptosis in creating and maintaining luminal space within normal and oncogene-expressing mammary acini. Cell, 2002. 111(1): p. 29–40.

29. Sicking, I., et al., Prognostic influence of cyclooxygenase-2 protein and mRNA expression in node-negative breast cancer patients. BMC Cancer, 2014. 14: p. 952.

30. Bernhardt, S.M., et al., Isogenic Mammary Models of Intraductal Carcinoma Reveal Progression to Invasiveness in the Absence of a Non-Obligatory In Situ Stage. Cancers (Basel), 2023. 15(8).

31. Fornetti, J., et al., Physiological COX-2 expression in breast epithelium associates with COX-2 levels in ductal carcinoma in situ and invasive breast cancer in young women. Am J Pathol, 2014. 184(4): p. 1219–1229.

32. Smith, R., et al., Using Microarrays to Interrogate Microenvironmental Impact on Cellular Phenotypes in Cancer. J Vis Exp, 2019(147).

33. Watson, S.S., et al., Microenvironment-Mediated Mechanisms of Resistance to HER2 Inhibitors Differ between HER2+ Breast Cancer Subtypes. Cell Syst, 2018. 6(3): p. 329–342 e6.

34. Roh, J.S. and D.H. Sohn, Damage-Associated Molecular Patterns in Inflammatory Diseases. Immune Netw, 2018. 18(4): p. e27.

35. Scher, J.U. and M.H. Pillinger, 15d-PGJ2: the anti-inflammatory prostaglandin? Clin Immunol, 2005. 114(2): p. 100–9.

36. Luond, F., et al., Distinct contributions of partial and full EMT to breast cancer malignancy. Dev Cell, 2021. 56(23): p. 3203–3221 e11.

37. Ryu, S., et al., The matricellular protein SPARC induces inflammatory interferon-response in macrophages during aging. Immunity, 2022. 55(9): p. 1609–1626 e7.

38. Lin, Y., S. Jamison, and W. Lin, Interferon-gamma activates nuclear factor-kappa B in oligodendrocytes through a process mediated by the unfolded protein response. PLoS One, 2012. 7(5): p. e36408.

39. Allen, J.E., IL-4 and IL-13: Regulators and Effectors of Wound Repair. Annu Rev Immunol, 2023. 41: p. 229–254.

40. Larson, C., et al., TGF-beta: a master immune regulator. Expert Opin Ther Targets, 2020. 24(5): p. 427–438.

41. McDaniel, S.M., et al., Remodeling of the mammary microenvironment after lactation promotes breast tumor cell metastasis. Am J Pathol, 2006. 168(2): p. 608–20.

42. Guo, Q., et al., Mammary collagen is under reproductive control with implications for breast cancer. Matrix Biol, 2022. 105: p. 104–126.

43. Esbona, K., et al., COX-2 modulates mammary tumor progression in response to collagen density. Breast Cancer Res, 2016. 18(1): p. 35.

44. Nagaharu, K., et al., Tenascin C induces epithelial-mesenchymal transition-like change accompanied by SRC activation and focal adhesion kinase phosphorylation in human breast cancer cells. Am J Pathol, 2011. 178(2): p. 754–63.

45. Xu, C., et al., SPP1, analyzed by bioinformatics methods, promotes the metastasis in colorectal cancer by activating EMT pathway. Biomed Pharmacother, 2017. 91: p. 1167–1177.

46. Liu, Y., et al., Pleiotropic Effects of PPARD Accelerate Colorectal Tumorigenesis, Progression, and Invasion. Cancer Res, 2019. 79(5): p. 954–969.

47. Friis, S., et al., Low-Dose Aspirin or Nonsteroidal Anti-inflammatory Drug Use and Colorectal Cancer Risk: A Population-Based, Case-Control Study. Ann Intern Med, 2015. 163(5): p. 347–55.

48. Basudhar, D., et al., Coexpression of NOS2 and COX2 accelerates tumor growth and reduces survival in estrogen receptor-negative breast cancer. Proc Natl Acad Sci U S A, 2017. 114(49): p. 13030–13035.

49. Cheng, R.Y.S., et al., Interferon-gamma is quintessential for NOS2 and COX2 expression in ER(-) breast tumors that lead to poor outcome. Cell Death Dis, 2023. 14(5): p. 319.

50. Zhang, L., M. Zeng, and B.M. Fu, Inhibition of endothelial nitric oxide synthase decreases breast cancer cell MDA-MB-231 adhesion to intact microvessels under physiological flows. Am J Physiol Heart Circ Physiol, 2016. 310(11): p. H1735–47.

51. Fahey, J.M. and A.W. Girotti, Nitric oxide-mediated resistance to photodynamic therapy in a human breast tumor xenograft model: Improved outcome with NOS2 inhibitors. Nitric Oxide, 2017. 62: p. 52–61.

52. Boyle, A.J., et al., Repurposing [(11)C]MC1 for PET Imaging of Cyclooxygenase-2 in Colorectal Cancer Xenograft Mouse Models. Mol Imaging Biol, 2022. 24(3): p. 365–370.

53. Grit, J.L., et al., Ex Vivo Patient-Derived Explant Model for Neurofibromatosis Type 1-Related Cutaneous Neurofibromas. J Invest Dermatol, 2024. 144(9): p. 2052–2065 e8.

54. O’Brien, J., et al., Non-steroidal anti-inflammatory drugs target the pro-tumorigenic extracellular matrix of the postpartum mammary gland. Int J Dev Biol, 2011. 55(7-9): p. 745–55.

