## Supplementary material for "S-Nitrosylated COX-2 is a TME-regulated breast cancer biomarker of mesenchymal phenotypes": Full Supplemental File

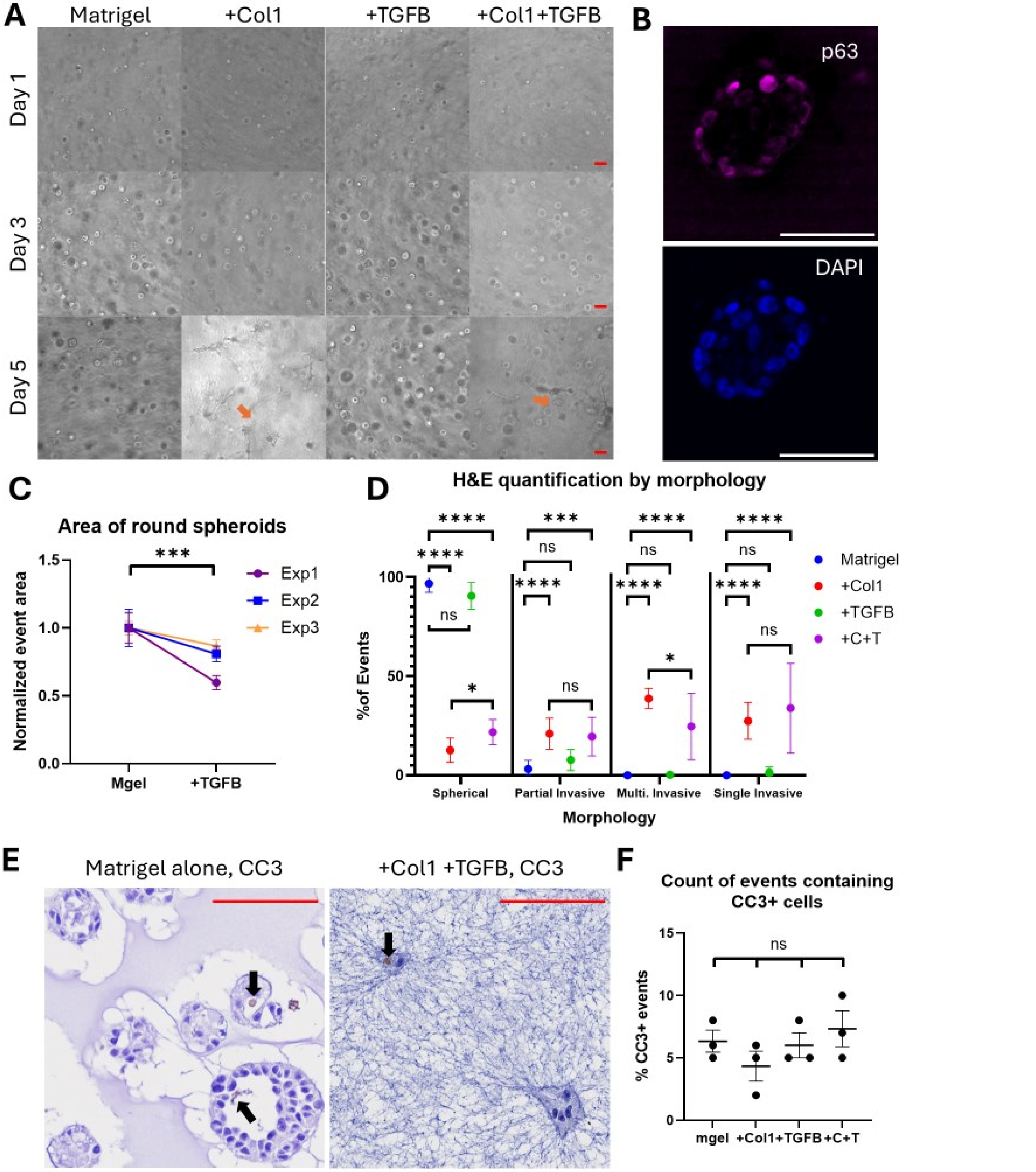

**Supplementary Figure 1: 3D cell culture time course, +TGFB spheroid size quantitation, H&E event count details, and CC3 staining. A)** Brightfield images of MCF10DCIS.com cells in 3D Matrigel culture, with or without the addition of fibrillar collagen 1 and TGFB1. Images were taken live on culture days 1, 3, and 5. Scale bars (red) are 100 µm. **B)** Representative image demonstrating myoepithelial biomarker p63 staining in “basal” cells when grown in matrigel alone, consistent with the bipotential nature of this cell line. Scale bars (white) are 50 µm. **C)** Area (size) of spheroids in matrigel-alone (mgel) compared to +TGFB conditions across three experiments, normalized to matrigel average. **D)** Expansion of Fig 1B; percentage of events of each phenotype in each condition from H&E counts with statistical comparison by multivariable modeling; points are averages, whiskers are standard error of the mean. **E)** Representative IHC staining images for cleaved caspase 3 (CC3) with arrows indicating rare positive cells. Scale bars (red) are 100 µm. **F)** Quantification of CC3 staining as the percentage of spheroids with at least one CC3+ cell in each condition in each experiment. Points indicate individual experimental averages, horizontal bold bar indicates average, whiskers show standard error of the mean. **All)** Statistical significance is shown as *, **, *** and **** for p values < 0.05, 0.01, 0.001, and 0.0001 respectively. “ns” or unlisted relationships were not significant.

| | SNAIL | Vimentin | N-cadherin | COX-2 | GAPDH |
|---|---|---|---|---|---|
| Matrigel | 31.07 ± 0.29 | 18.97 ± 0.23 | 32.28 ± 0.39 | 29.74 ± 0.35 | 17.79 ± 0.15 |
| +Col1 | 31.95 ± 0.21 | 20.06 ± 0.13 | 32.13 ± 0.19 | 26.17 ± 0.34 | 18.61 ± 0.12 |
| +TGFB | 28.79 ± 0.31 | 18.88 ± 0.18 | 27.84 ± 0.54 | 26.70 ± 0.36 | 18.77 ± 0.16 |
| +Col1+TGFB | 27.92 ± 0.35 | 19.02 ± 0.24 | 27.19 ± 0.36 | 25.00 ± 0.20 | 18.88 ± 0.19 |
| Overall | 29.99 ± 0.34 | 19.24 ± 0.14 | 29.91 ± 0.49 | 26.96 ± 0.37 | 18.50 ± 0.11 |

**Supplementary Table 1: Non-normalized, average Cq values for each RT-qPCR primer across experiments.** Cq values listed are averaged from three qPCR replicates of three technical replicates of three experimental replicates within each condition or in grand total (overall), presented with a 95% confidence interval.

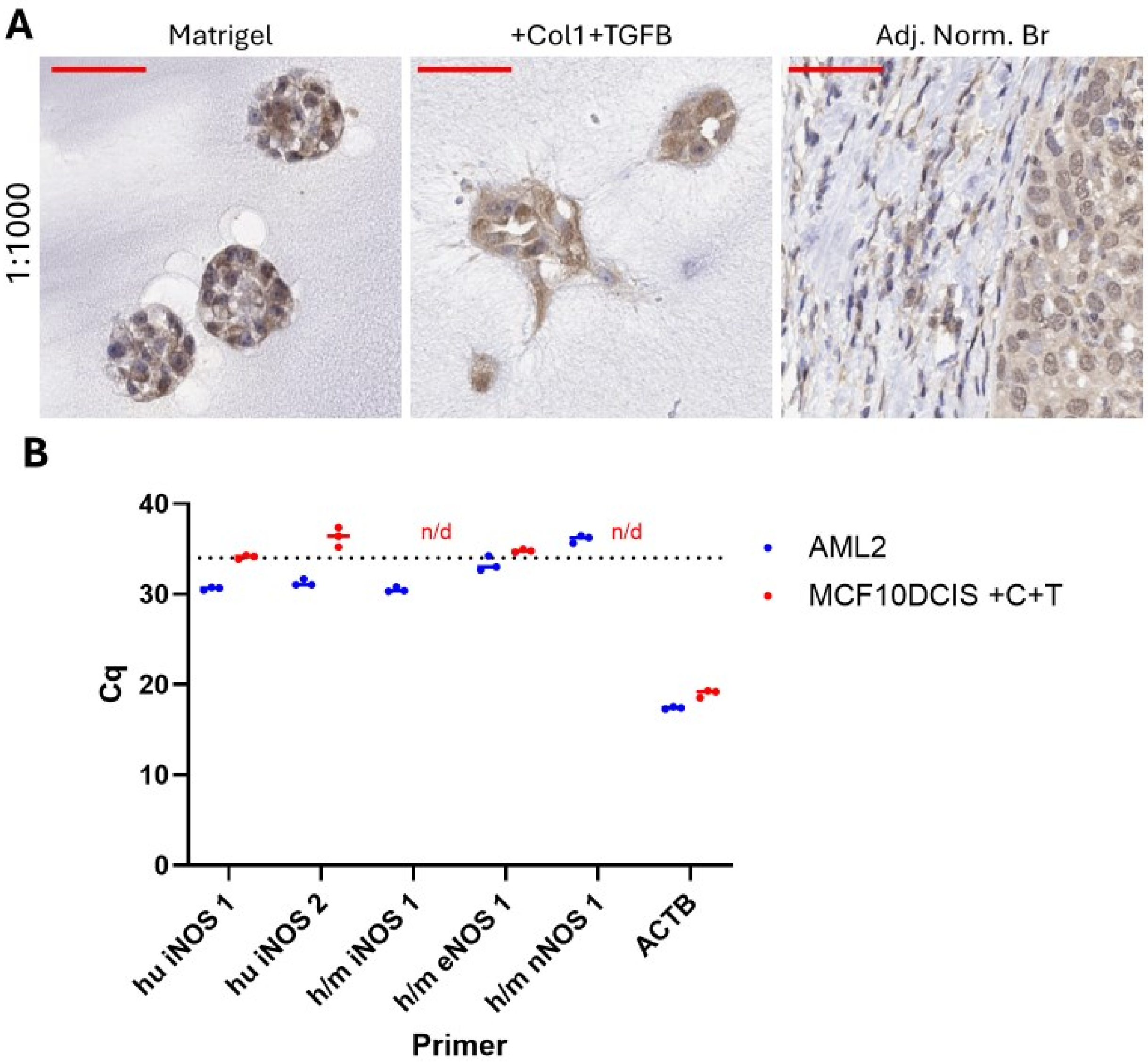

**Supplementary Figure 2: Human Vimentin staining and NOS RT-qPCR. A)** Staining using anti-Vimentin primary antibodies at 1:1000 concentration on MCF10DCIS in two 3D culture conditions (matrigel alone, +Col+TGFB) and adjacent normal human breast tissue as positive and negative controls. Stain is brown, nuclear counter stain is blue. Overall vimentin staining in MCF10DCIS.com cells was high. Scale bars = 50 μm. **B)** RT-qPCR assay using primers for iNOS, eNOS, or nNOS; primers are designed for human NOS mRNA (hu) or to be compatible with both human and mouse mRNA (h/m). Actin Beta (ACTB) is used as a housekeeping gene for loading control. Lysate from human acute myeloid leukemia AML2 cell line was used as a positive mRNA control for iNOS expression. The experimental sample is from MCF10DCIS in +Col+TGFB 3D culture. Data is shown as raw Cq values of a single sample run in triplicate; bars indicate average Cq, n/d indicates the fluorescent threshold was never reached. A dotted line at Cq = 34 indicates the point above which results become unreliable.

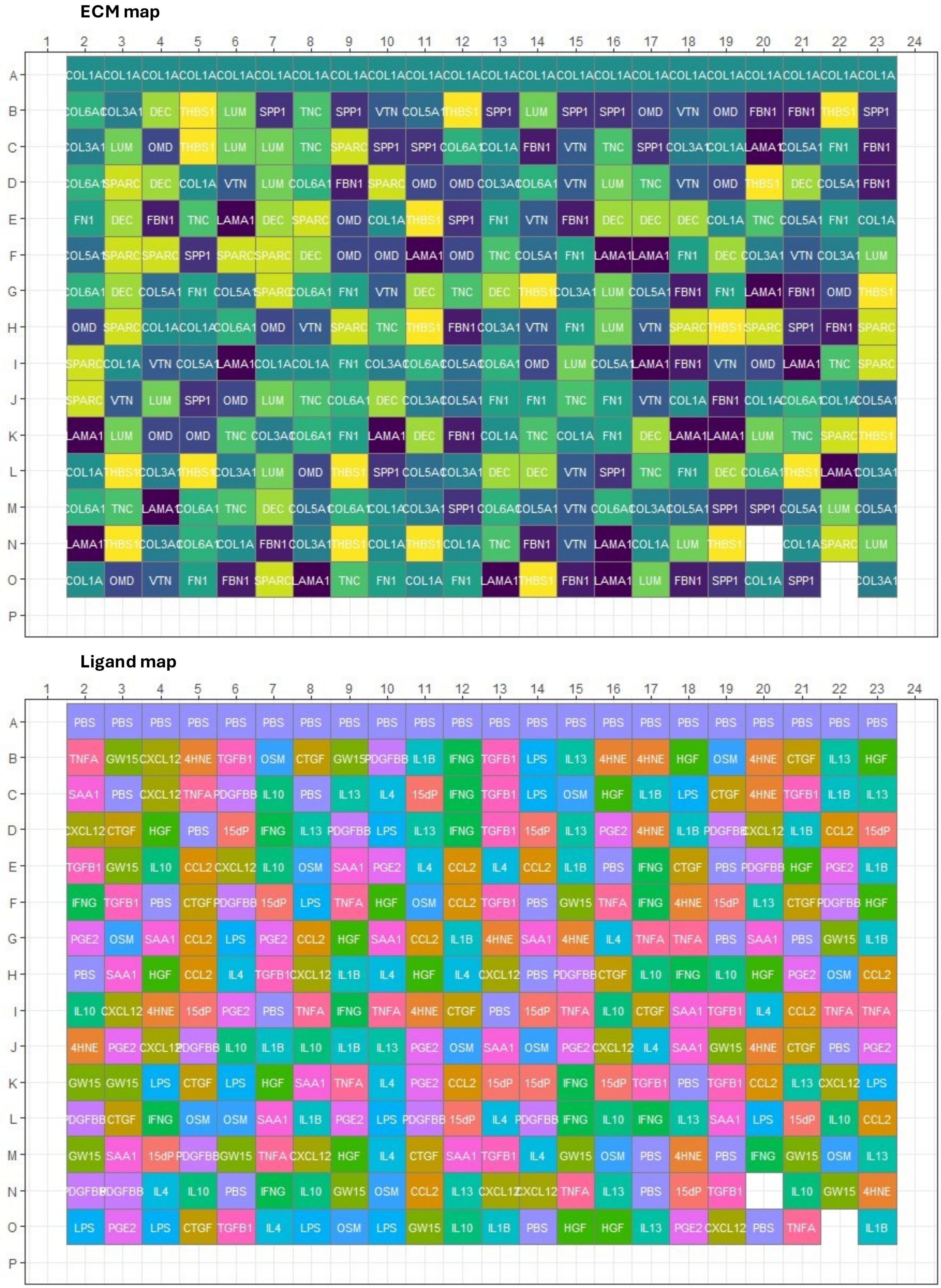

**Supplementary Figure 3: MEMA plate ECM and ligand arrangement, cell count, and intensity distribution.**

Plate map for ECM (top) and ligand (bottom) combinations used in MEMA assay. Row A wells were seeded with HCA-7 cells as a positive control for SNO-COX-2, rows B-O were seeded with MCF10DCIS cells. Wells N20 and O22 received no ECM coating and were seeded as a negative plating control.

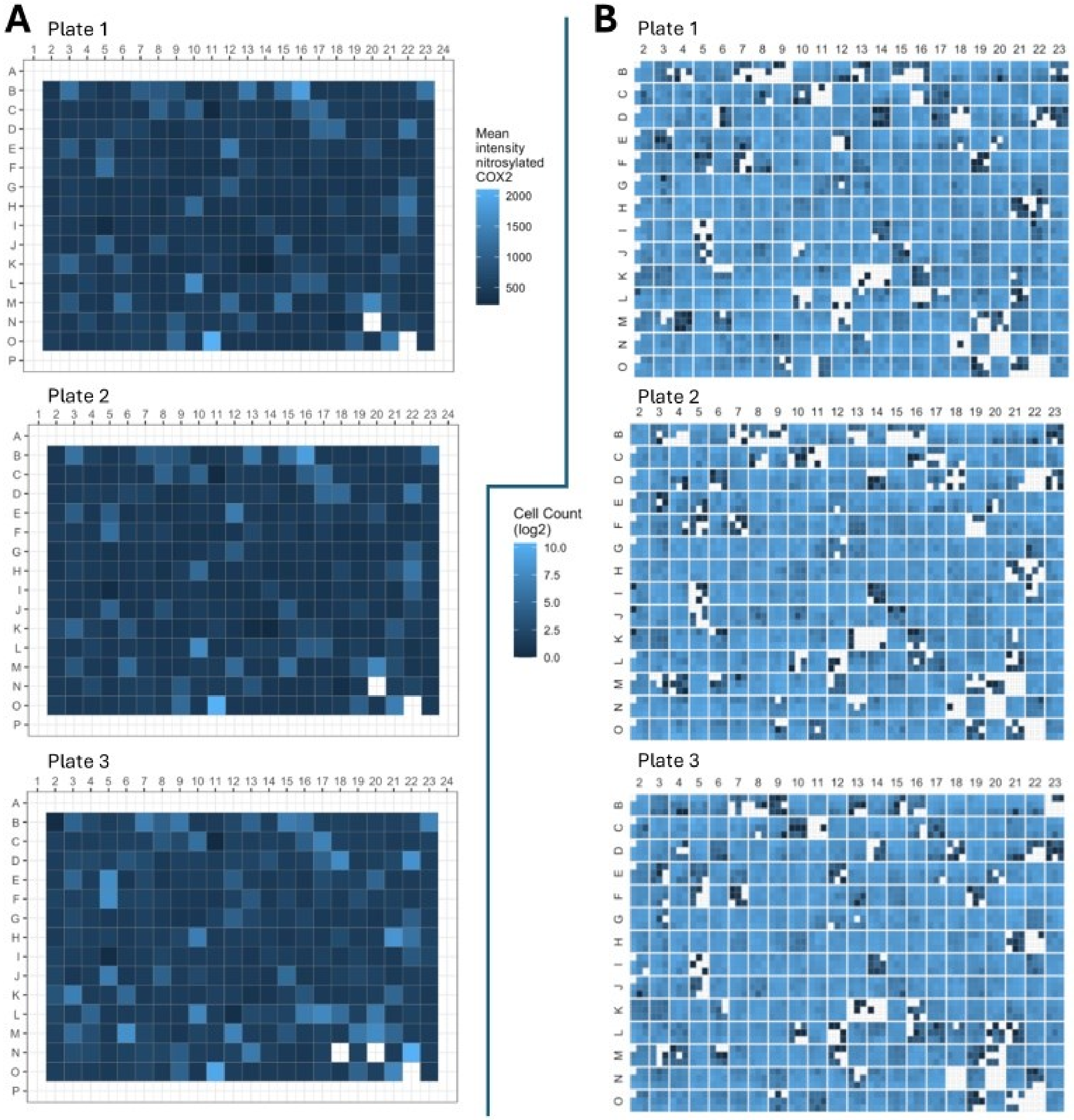

**Supplementary Figure 4: MEMA cell counts and intensity distribution. A)** Mean SP21 intensity per cell per well across images in each MEMA plate. **B)** Mean cell count per image (right, images in 3x3 grid within each well); white squares indicate no cells present.

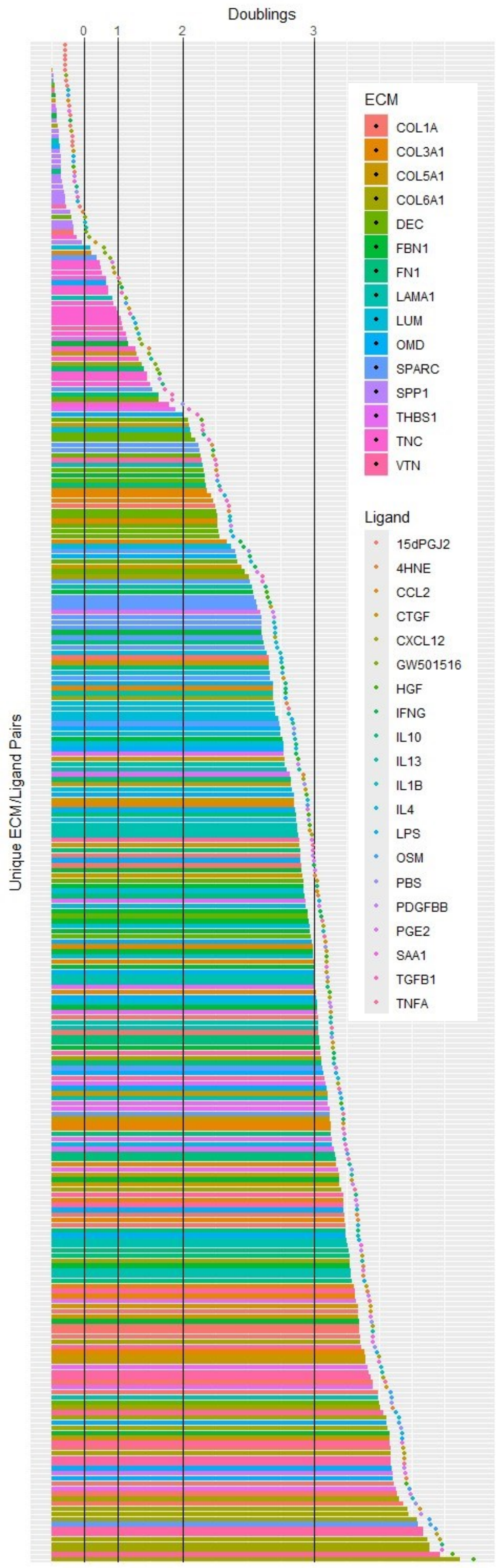

**Supplementary Figure 5: Average cell counts per condition in the MEMA assay in ascending order.** Horizontal bars indicate average cell count per image per condition. Bars are colored to indicate ECM factors of each condition; lifted dots are colored to indicate the paired ligand of each. Black lines indicate roughly 0, 1, 2, and 3 doublings; calculations in methods. Note that dot position are displaced relative to their matched bars for visualization and do not represent doublings.

| ECM factor | Names | Supplier | Cat# | Conc. |
|---|---|---|---|---|
| COL1A | Collagen type I alpha 1 chain | Cultrex | 3442-050-01 | 1 μg/cm$^2$ |
| COL3A1 | Collagen type III alpha 1 chain | Millipore | CC054 | 1 μg/cm$^2$ |
| COL5A1 | Collagen Type V Alpha 1 Chain | Millipore | CC077 | 1 μg/cm$^2$ |
| COL6A1 | Collagen Type VI Alpha 1 Chain | COLVI | 354261 | 1 μg/cm$^2$ |
| DEC | Decorin; **DCN** | R&D Systems | 143-DE | 1 μg/cm$^2$ |
| FBN1 | Fibrillin 1 | Lynn Sakai Lab | N/A | 1 μg/cm$^2$ |
| FN1 | Fibronectin 1 | R&D Systems | 1918-FN | 1 μg/cm$^2$ |
| LAMA1 | Laminin Subunit Alpha 1 | Sigma | L6274 | 1 μg/cm$^2$ |
| LUM | Lumican | R&D Systems | 2846-LU | 1 μg/cm$^2$ |
| OMD | Osteomodulin | R&D Systems | 2884-AD | 1 μg/cm$^2$ |
| SPARC | Secreted Protein Acidic And Cysteine Rich; Osteonectin | R&D Systems | 941-SP | 1 μg/cm$^2$ |
| SPP1 | Secreted phosphoprotein 1, **Osteopontin** | R&D Systems | 1433-OP | 1 μg/cm$^2$ |
| THBS1 | Thrombospondin 1 | R&D Systems | 3074-TH | 1 μg/cm$^2$ |
| TNC | Tenascin C | R&D Systems | 3358-TC | 1 μg/cm$^2$ |
| VTN | Vitronectin | R&D Systems | 2308-VN | 1 μg/cm$^2$ |
| **Ligand** | **Names** | **Supplier** | **Cat#** | **Conc.** |
| 15dPGJ2 | 5-Deoxy-δ12,14-Prostaglandin J2 | Cayman | 18570 | 10 ug/mL |
| 4HNE | 4-Hydroxynonenal | Cayman | 32100 | 1 μM |
| CCL2 | C-C Motif Chemokine Ligand 2; MCP1 | R&D Systems | 279-MC | 0.03 ug/mL |
| CTGF | Connective Tissue Growth Factor; **CCN2** | Life Tech | PHG0286 | 0.05 μM |
| CXCL12 | C-X-C Motif Chemokine Ligand 12; PBSF | R&D Systems | 350-NS-010 | 0.01 ug/mL |
| GW501516 | Cardarine | Selleck | S5615 | 50 μM |
| HGF | Hepatocyte Growth Factor | R&D Systems | 294-HG | 0.04 ug/mL |
| IFNG | Interferon Gamma | R&D Systems | 285-IF | 0.02 ug/mL |
| IL10 | Interleukin 10 | R&D Systems | 1064-IL | 0.01 ug/mL |
| IL13 | Interleukin 13 | R&D Systems | 213-ILB | 0.01 ug/mL |
| IL1B | Interleukin 1 Beta | R&D Systems | 201-LB | 0.001 ug/mL |
| IL4 | Interleukin 4 | R&D Systems | 6507-IL-010 | 0.01 ug/mL |
| LPS | Lipopolysaccharide; endotoxin | Sigma | L2654 | 0.005 ug/mL |
| OSM | Oncostatin M | R&D Systems | 8475-OM | 0.01 ug/mL |
| PBS | Phosphate Buffered Saline (control) | | | |
| PDGFBB | Platelet Derived Growth Factor Subunit B; PDGF2 | R&D Systems | 220-BB | 0.05 ug/mL |
| PGE2 | Prostaglandin E2 | Sigma | P6532-1MG | 0.001 ug/mL |
| SAA1 | Serum Amyloid A1 | Peprotech | AF-300-53 | 1 ug/mL |
| TGFB1 | Transforming Growth Factor Beta 1 | R&D Systems | 7754-BH | 0.01 ug/mL |
| TNFA | Tumor Necrosis Factor | R&D Systems | 210-TA | 0.01 ug/mL |

**Supplementary Table 2: MEMA ECM and Ligand details.** Includes purchasing information and the final concentration (Conc.) per well used in the MEMA assay. Note that ECM factor concentration is measured against the area of the plate well they cover.

| mRNA Target | Species | Forward Primer Sequence | Reverse Primer Sequence | Product Length | Note |
|---|---|---|---|---|---|
| SNAIL | human | CAGTGCCTCGACCACTAT | CTGCTGGAAGGTAAACTCTGG | 115 | |
| N-Cadherin | human | GAGGAGTCAGTGAAGGAGTC | GGAGTTTTCTGGCAAGTTGA | 132 | |
| Vimentin | human | ATGTTGACAATGCGTCTCTG | CTCCTGGATTTCCTCTTCGT | 107 | |
| COX-2 | human | CTGGGAAGCCTTCTCTAACC | GGAAGCTGCTTTTTACCTTTG | 102 | |
| GAPDH | human | TCAAGATCATCAGCAATGCC | ATGAGTCCTTCCACGATACC | 94 | Housekeeper |
| iNOS | human | CATCTACCAGGAGGAGATGC | TCCTGAACATAGACCTTGGG | 105 | Minor trial |
| iNOS | human | GGAGACGGGAAAGAAGTCTC | CCCCAGTTTTTGATCCTCAC | 90 | Minor trial |
| iNOS | hu/mus | CTCCATAGTTTCCAGAAGCA | GGAGGGACCAGCCAAATC | 134 | Minor trial |
| eNOS | hu/mus | CCCTCACCGCTACAACAT | CTGCCTTGTCTTTCCACAG | 89 | Minor trial |
| nNOS | hu/mus | CGCCTTAGATCTGAGTCCAT | GTTAGGAGCTGAAAACCTCAT | 73 | Minor trial |
| ACTB | human | CCTGTACGCCAACACAGTGC | ATACTCCTGCTTGCTGATCC | 211 | Housekeeper |

**Supplementary Table 3: RT-qPCR primer list.** Species indicates whether primers were designed for human mRNA (hu) or to be compatible with both human and mouse orthologues of the same gene (mu). All sequences are given 5' to 3'. Product length is in nucleotides.
